# Dissecting the import and export pathways of the human RNA helicase UPF1

**DOI:** 10.1101/2021.08.02.454737

**Authors:** Andrea B. Eberle, Karin Schranz, Sofia Nasif, Lena Grollmus, Oliver Mühlemann

## Abstract

The RNA helicase UPF1 is best known for its key role in mRNA surveillance but has been implicated in additional cellular processes both in the nucleus and in the cytoplasm. In human cells, the vast majority of UPF1 resides in the cytoplasm and only small amounts can be detected in the nucleus at steady state. It was previously shown that its export from the nucleus to the cytoplasm is Crm1-dependent, yet neither the nuclear export signal (NES) nor the nuclear localization signal (NLS) has been identified. Here, we provide evidence for a noncanonical NLS in UPF1, map the NES to amino acids 89-105 and show that L103 and F105 are essential for UPF1’s export to the cytoplasm. Examination of additional UPF1 mutants revealed that a functional helicase domain but not the association with RNA is crucial for the shuttling capacity of UPF1.

## Introduction

From its transcription in the cell nucleus to its translation in the cytoplasm messenger ribonucleoparticles (mRNPs), which consist of mRNA and all proteins bound to it, undergo constant remodelling with various RNA-binding proteins (RBPs) leaving and joining the mRNP (Singh et al., 2015). These remodelling events alter the overall structure of the mRNP and thereby have important regulatory impact on mRNA synthesis, processing, localization, translation, and stability, and hence on the expression of the genetic information. RNA helicases play key roles in these energy-dependent mRNP remodelling steps and therefore exert crucial functions in post-transcriptional gene expression (Jankowsky, 2011).

Up-frameshift 1 (UPF1) is an abundant ATP-dependent RNA helicase belonging to the non-ring forming superfamily (SF) 1 of eukaryotic RNA helicases, which contain a conserved helicase core that consists of two RecA-like domains. The two RecA-like domains form the ATP-binding pocket and a composite RNA-binding surface (Chakrabarti et al., 2011; Cheng et al., 2007; Gowravaram et al., 2018). Driven by ATP hydrolysis, UPF1 unwinds both RNA and DNA in a highly processive manner and translocates on nucleic acid in 5’ to 3’direction (Fiorini et al., 2015). The helicase domain (HD) of UPF1 is flanked N-terminally by a cysteine-histidine rich (CH) domain and at the C-terminus by an unstructured stretch of amino acids enriched in serine-glutamine (SQ) motifs (Figure 1A), both of which were shown to suppress the catalytic activity of UPF1 (Chamieh et al., 2008; Fiorini et al., 2013).

**Figure 1.**
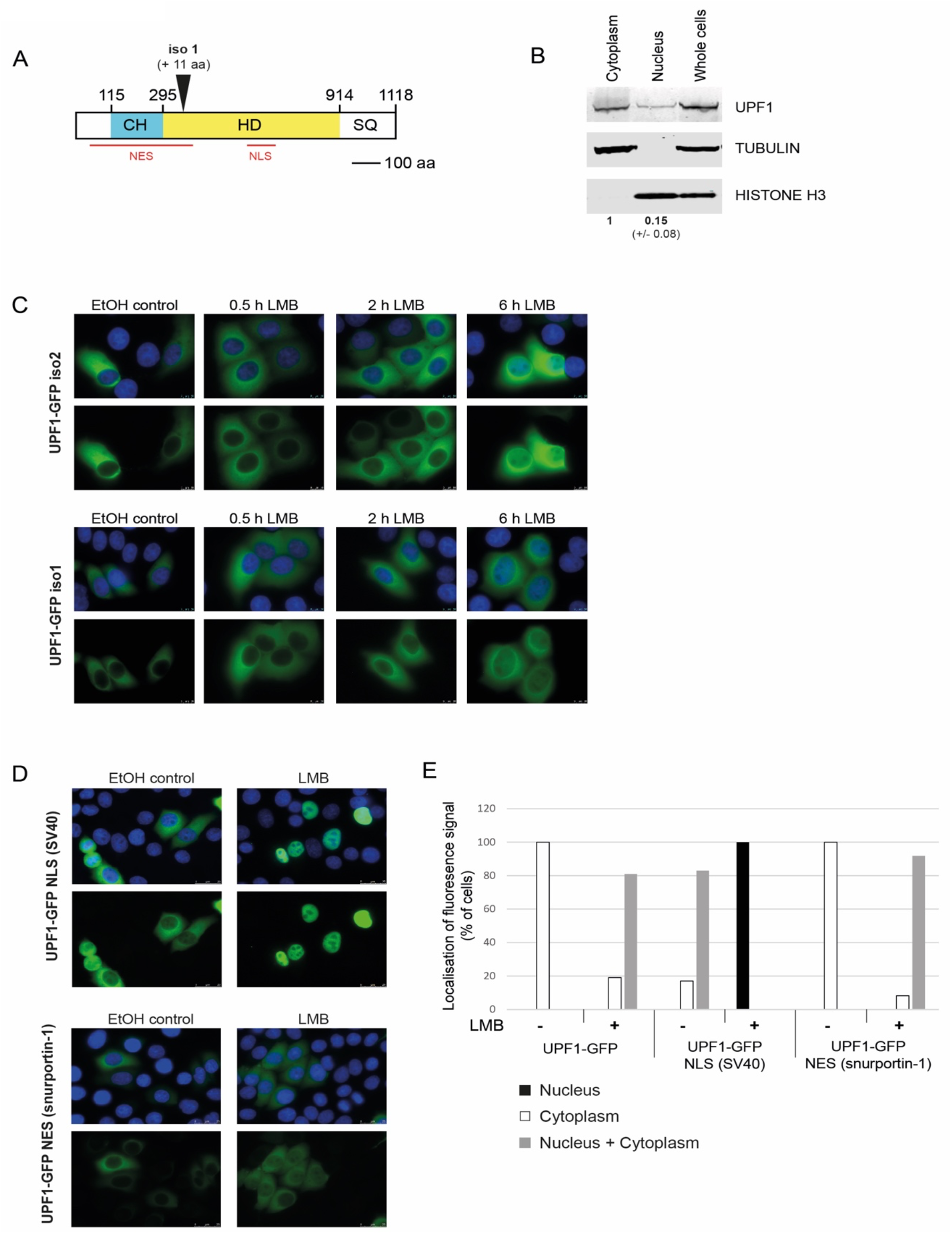
UPF1 is a shuttling protein. **A**. Schematic structure of UPF1 including NLS and NES postulated by Mendell [adapted from (Kim and Maquat, 2019; Mendell *et al*., 2002)]. **B**. Localization of UPF1 in HeLa cells was assessed by Western Blotting of nuclear and cytoplasmic fractions. Detection of TUBULIN and HISTONE H3 was used to control the quality of the biochemical fractionations. Quantification by Image Studio software of the UPF1 signal in three independent experiments showed a ratio of nucleus to cytoplasm of 0.15 (+/- 0.08). 2 x 10^5^ cell equivalents were loaded in each lane. **C**. Plasmids encoding UPF1-GFP isoform 2 (iso2) or isoform 1 (iso1) were transfected into HeLa cells and 48 hours later analyzed by fluorescence microscopy when no LMB was added (EtOH control) or when LMB was added for 0.5, 2 or 6 hours. Pictures of the GFP signal in the green channel and a composite of the green and blue channel (DAPI staining for nucleus) are shown. **D**. NLS of SV40 (PKKKRKV) or NES of snurportin-1 (EELSQALASSFSV) was attached at the C-terminus of UPF-GFP. Transfected HeLa cells treated without LMB or with LMB (6 hours) were examined by fluorescence microscopy as in C. **E**. GFP signals of individual cells (from experiments in B and C) were observed and classified into complete nuclear localization (black bars), complete cytoplasmic localization (white bars) or localization in nucleus and cytoplasm (grey bars). Cells of two independent experiments were analyzed: UPF1-GFP n=68/105 (-LMB/+LMB); UPF1-GFP NLS (SV40) n=47/55 (-LMB/+LMB); UPF1-GFP NES (snurportin-1) n=78/61 (-LMB/+LMB).

With about 360’000 molecules per cell, human cells contain roughly the same amount of UPF1 molecules as mRNA molecules (Hein et al., 2015). UPF1 is an essential protein in mammalian cells: UPF1 knockout is embryonic lethal (Medghalchi et al., 2001) and immortalized cells stop dividing and die about 5-7 days after UPF1 depletion (Azzalin and Lingner, 2006). UPF1 has been implicated in several different cellular processes among which its key role in nonsense-mediated mRNA decay (NMD) is best characterized (Karousis and Muhlemann, 2019; Kim and Maquat, 2019). Besides its essential role in NMD (see below), UPF1 has been associated with additional mRNA decay pathways that are specified by different UPF1-interacting RBPs, such as Staufen [STAU; (Kim et al., 2005; Park and Maquat, 2013)], stemloop-binding protein [SLBP; (Choe et al., 2014; Kaygun and Marzluff, 2005)], glucocorticoid receptor [GR; (Cho et al., 2015; Park et al., 2016)], regnase 1 (Mino et al., 2015), and in tudor staphylococcal/micrococcal-like nuclease (TSN)-mediated microRNA decay (Elbarbary et al., 2017).

NMD serves a dual role as a mRNA quality control mechanism that targets for degradation aberrant mRNAs with a premature termination codon (PTC), which most frequently arise due to the inclusion or exclusion of an alternative exons (Karousis et al., 2021), and as a translation-dependent post-transcriptional regulator of mRNA stability for a subset of mRNAs that encode full length functional proteins (Karousis and Muhlemann, 2019; Kurosaki et al., 2019). The molecular mechanism of NMD, in particular the step of target mRNA selection, is still incompletely understood but it is well established that UPF1 is essential for NMD in all eukaryotes (Karousis and Muhlemann, 2019; Kurosaki *et al*., 2019). Cross-linking followed by immunoprecipitation (CLIP) studies detected UPF1 binding to essentially all mRNAs at many different sites over their entire length before the onset of translation, followed by a translation-dependent displacement from coding sequences, resulting in an enrichment on 3’ untranslated regions (UTRs) at cellular steady state (Hurt et al., 2013; Zund et al., 2013). On mRNAs on which ribosomes fail to terminate translation correctly (Karousis and Muhlemann, 2019), SMG1-mediated phosphorylation of serine and threonine of the serine/glutamine (SQ) and threonine/glutamine (TQ) motifs in the N- and C-terminal parts of UPF1 is observed (Durand et al., 2016; Kurosaki and Maquat, 2013; Lee et al., 2015; Yamashita et al., 2001), which appears to be the activating step in NMD. Hyperphosphorylated UPF1 then recruits the downstream effectors SMG5, SMG6 and SMG7, of which SMG6 is an endonuclease that induces the degradation of the targeted mRNA by cleaving it in the vicinity of the termination codon (Boehm et al., 2021; Eberle et al., 2009; Huntzinger et al., 2008; Lykke-Andersen et al., 2014).

Moreover, there is also evidence that UPF1 exerts different functions in the nucleus that contribute to genome stability (DNA damage response and R-loop formation) and the regulation of S-phase progression (Azzalin and Lingner, 2006; Dehghani-Tafti and Sanders, 2017; Ngo et al., 2021), as well as to the regulation of telomeric repeat-containing RNA (TERRA) and telomere replication (Azzalin et al., 2007; Chawla et al., 2011). In addition, UPF1 has been implicated in the nuclear export of HIV-1 genomic RNA (Ajamian et al., 2015) and data from *Drosophila* cells indicates that UPF1 associates with mRNAs co-transcriptionally and that it is required for their release from the gene loci (Singh et al., 2019).

Although most UPF1 is found in the cytoplasm of human cells under physiological conditions, in accordance with its translation-dependent function in NMD, a fraction of UPF1 can be detected in the nucleus of human cells (Ajamian *et al*., 2015; Hong et al., 2019; Mendell et al., 2002). In fact, UPF1 appears to be a shuttling protein and nuclear accumulation can be triggered by addition of leptomycin B (LMB) (Mendell *et al*., 2002), a *Streptomyces* metabolite that blocks CRM1 (Exportin 1)-dependent nuclear export by covalently binding to cysteine 528 in the nuclear export signal (NES)-binding region of CRM1 (Rahmani and Dean, 2017). Most proteins that depend on active import into the nucleus via the nuclear pore complex possess a nuclear localization signal (NLS) that binds to an import receptor of the importin superfamily. NLSs can consist of very different amino acid sequences but they are usually rich in positively charged residues (Lu et al., 2021). Classical monopartite NLSs bind to the major binding pocket of the adaptor protein importin α, while the noncanonical NLSs classes bind the minor binding pocket of importin α (Kosugi et al., 2009a). Importin α then interacts with importin β (aka karyopherin β1), which in turn contacts the nuclear pore complex and facilitates translocation of the cargo/importin complex into the nucleus (Lott and Cingolani, 2011). Alternatively, shuttling proteins can be directly bound by importin β independent of an adaptor protein (Lee et al., 2006), which makes computational predictions of NLSs challenging.

Here, we set out to functionally delineate the NLS and NES of UPF1. We confirmed that a previously reported large stretch of about 100 amino acids in the helicase domain of UPF1 is required for its import into the nucleus (Mendell *et al*., 2002) but were unable to further map this putative NLS, suggesting that UPF1 does not contain a NLS that resembles any of the known classes of NLSs (Kosugi *et al*., 2009a; Lu *et al*., 2021). However, we were able to precisely map the NES of UPF1 and identified two amino acids that when mutated inactivate the NES, resulting in accumulation of UPF1 in the nucleus. By studying a UPF1 mutant that is unable to bind RNA, we further revealed that the capability of binding RNA is not required for the import and export of UPF1, however, functional helicase activity is crucial for its nuclear import.

## Results

### Both UPF1 isoforms are shuttling proteins localizing mainly to the cytoplasm at steady state

UPF1 has been reported to shuttle between the nucleus and the cytoplasm, with the bulk part of the protein localizing to the cytoplasm at steady state in human and Drosophila cells (Lykke-Andersen et al., 2000; Mendell *et al*., 2002; Singh *et al*., 2019). In biochemical fractionation experiments of HeLa cells, we detected about 7 times less UPF1 in the nuclear fraction than in the cytoplasmic fraction (Figure 1B). Shuttling proteins typically contain defined amino acids sequences functioning as nuclear localization signal (NLS) and nuclear export signal (NES). A rough characterization of the NLS and NES in UPF1, undertaken almost 20 years ago, indicated that the NES resides in the N-terminal part of the protein somewhere between amino acids 55 and 416, and the NLS lies within the helicase domain between amino acids 596 and 697 (Figure 1A), since deletions of these regions led to an accumulation of UPF1 in the nucleus or its failure to localize to the nucleus, respectively (Mendell *et al*., 2002). However, in this first characterization large deletions of 100 amino acids or more were used to impair the shuttling properties of UPF1, which bears the risk of inducing conformational changes in the protein that indirectly affect nucleo-cytoplasmic transport. A more detailed mapping of the NLS and NES would be very useful for studying the reported functions of UPF1 in the nucleus and in the cytoplasm and clearly distinguish them from each other.

First, we addressed whether the two UPF1 isoforms found in human cells might differ in their subcellular localization. UPF1 isoform 2, which results from alternative pre-mRNA splicing and lacks 11 amino acids in a regulatory loop of the helicase domain, is more abundant in human cells and better characterized than isoform 1 (Gowravaram *et al*., 2018; Nicholson et al., 2014). The extended regulatory loop of UPF1 isoform 1 was found to increase UPF1’s translocation and ATPase activities by two-fold (Gowravaram *et al*., 2018), but apart from this no functional differences between the two isoforms are known. To investigate the localization of both UPF1 isoforms, enhanced green fluorescence protein (GFP) was fused via a linker of 17 glycine residues (G-linker) to the C-terminus of UPF1 isoform 1 (iso1) or isoform 2 (iso2). Similar expression of both GFP-tagged UPF1 isoforms in HeLa cells upon transient transfection of the expression plasmids was confirmed by Western blotting (Figure S1A). Examination of the cells under the fluorescence microscope revealed that both UPF1 isoforms were cytoplasmic at steady state and without any treatment (EtOH control; Figure 1C). For both isoforms, a weak nuclear accumulation of UPF1-GFP could be detected 2 hours after addition of leptomycin B (LMB) and a clear nuclear signal after 6 hours (Figure 1C). LMB specifically inhibits the nuclear export receptor CRM1 and therefore causes nuclear accumulation of shuttling proteins exported via the CRM1 pathway (Kudo et al., 1998). Since we detected no difference in localization between the two UPF1 isoforms, the predominant and better studied isoform 2 was used for all subsequent experiments.

To get insights into the shuttling properties of UPF1, a strong NLS from SV40 or a strong NES from snurportin-1 was attached to the C-terminus of UPF1-GFP (Kazgan et al., 2010) (Figures 1D and S1B). The addition of the strong SV40 NLS led to a more pronounced nuclear localization compared to UPF1-GFP, even in the absence of LMB, indicating that UPF1 intrinsic NLS is considerably weaker than the SV40 NLS (Figure 1C-E). However, since cytoplasmic GFP signal could be detected in all cells expressing UPF1-GFP NLS (SV40), there is evidence for a potent endogenous NES in UPF1. In the presence of LMB, the GFP signal was completely nuclear for UPF1-GFP NLS (SV40) in all cells observed (Figure 1D and E). Attaching the strong NES from snurportin-1 to the C-terminus of UPF1-GFP resulted in cytoplasmic localization in the absence of LMB and to cytoplasmic and nuclear GFP signal upon addition of LMB in about 90 % of the cells, proving that the UPF1-GFP NES (snurportin-1) protein was indeed shuttling and accumulated in the nucleus when export was impaired (Figure 1D and 1E). In contrast to the addition of a NLS, insertion of the snurportin-1 NES did not affect the localization behaviour of UPF1-GFP, further indicating that UPF1 possesses a potent endogenous NES with a comparable activity to the snurportin-1 NES (Figure 1C-E).

### No classical NLS is present in UPF1

Next, we attempted to define the NLS of UPF1 and inactivate it by point mutations to obtain an exclusively cytoplasmic version of UPF1. The bioinformatic tool *cNLS mapper* has been developed to predict importin α-dependent NLS (Kosugi et al., 2009b). Interestingly, the only predicted monopartite classical NLS (cNLS) in the UPF1 protein (score above 5.0) resided at amino acid positions 596-606 (RALKRTAEREL), right at the start of the putative 102 amino acids long NLS mapped previously (Mendell *et al*., 2002). Furthermore, no bipartite cNLS was predicted by *cNLS mapper*. The sequence encoding the putative NLS was deleted by fusion PCR from the UPF1-GFP construct, giving raise to the construct UPF1-GFP Δ596-606. In addition, a control plasmid in which the large NLS region proposed by Mendell was deleted was cloned (UPF1-G16-GFP Δ596-697). The plasmids were transfected into HeLa cells, the expression of the proteins was confirmed by Western blotting (Figure S1C) and the GFP signal was analyzed by fluorescence microscopy (Figure 2A). Confirming the initial study (Mendell *et al*., 2002), UPF1-GFP lacking amino acid residues 596-697 was entirely cytoplasmic even upon addition of LMB, indicating that the NLS is located in this region of the helicase domain (Figure 2A). In contrast, the deletion of the predicted NLS (amino acids 596-606) localized indistinguishable to wild-type (wt) UPF1-GFP with mainly cytoplasmic signal in the absence of LMB and a clear nuclear accumulation upon LMB addition, indicating that deletion of these 11 amino acids was not sufficient to impair protein import (Figure 2A).

**Figure 2.**
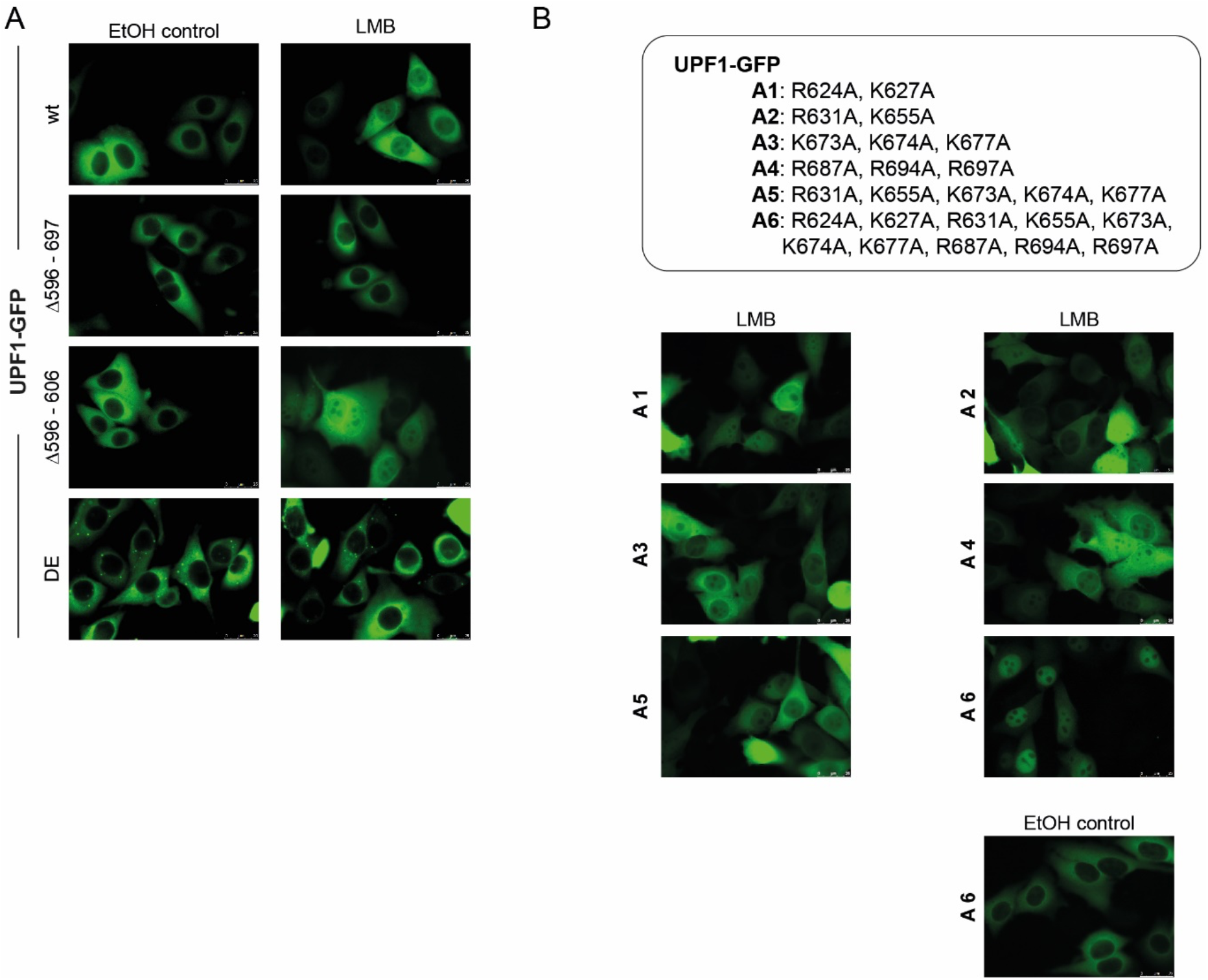
Investigation of UPF1’s NLS. **A**. Plasmids encoding UPF1-GFP wild type or different variants of UPF1 [deletions: UPF1-GFP Δ596-697 and UPF1-GFP Δ596-606; point mutations: D636A + E637A (DE)] were transfected into HeLa cells and the fluorescence signals were analyzed 48 hours after transfection in the control condition (EtOH control) and upon addition of LMB for 6 hours (LMB). **B**. Positively charged amino acids were replaced by alanines as explicated in the legend above the fluorescence pictures. Experiment was performed as in A. The EtOH control is shown only for UPF1-GFP A6.

For the classical protein import pathway, basic amino acids are typically crucial to allow for binding of importin α, which acts as an adapter protein between the NLS-containing cargo protein and importin β, which is responsible for the transport through the nuclear pore (Kosugi *et al*., 2009a). To test if UPF1 is transported to the nucleus by the importin α*/*β pathway, all 10 positively charged amino acids in the region 607-697 were replaced by alanines (all together (A6) or in different combinations (A1-A5); Figures 2B and S1C). If two to five lysines/arginines were replaced (constructs A1-A5), UPF1-GFP behaved like wt with cytoplasmic localization in control conditions and nuclear and cytoplasmic signal after treatment with LMB (Figure 2B). When all 10 positively charged amino acids were changed (A6), the GFP signal was also cytoplasmic in the absence of LMB as for the other constructs, however and unexpectedly, an almost exclusively nuclear signal was detected in the presence of LMB. To further analyze this curious finding, we identified by mass spectrometry proteins that co-precipitated with the A6 UPF1-GFP mutant and found many heat shock proteins, suggesting that this mutant fails to fold properly (Figure S1D). Collectively from this series of experiments attempting to decipher the NLS of UPF1, we conclude that UPF1 most likely does not contain a classical NLS and that it is imported into the nucleus in an importin α–independent manner.

### Helicase activity of UPF1 but not its binding to RNA is important for shuttling capacity

Interestingly, the highly conserved and well characterized amino acids aspartate and glutamate at amino acid positions 636 and 637 lie within the helicase region that is important for UPF1 import to the nucleus (Bhattacharya et al., 2000). When these two residues are mutated (DE variant), RNA and ATP can still bind to UPF1, but ATP hydrolysis and thus helicase activity is impaired, leading to P-body formation when this mutant is overexpressed in cells (Cheng *et al*., 2007; Franks et al., 2010) (Figure 2A). The UPF1-GFP DE mutant showed cytoplasmic localization even in the presence of LMB, suggesting that the helicase activity is necessary for moving to the nucleus (Figure 2A). Several studies indicated that this mutant is stuck on RNA, since dissociation from RNA requires ATP hydrolysis (Franks *et al*., 2010; Lee *et al*., 2015). This failure to dissociate from RNA leads to the formation of large RNA-protein aggregates (i.e. P-bodies), which we reasoned might sequester UPF1 in the cytoplasm and prevent its transport into the nucleus.

Therefore, we decided to test whether RNA binding is important for UPF1’s ability to shuttle between the nucleus and the cytoplasm. Based on the UPF1 crystal structure (Chakrabarti *et al*., 2011), it was predicted that the three amino acids asparagine 524, lysine 547 and arginine 843 might be engaged in and required for RNA binding (Sutapa Chakrabarti, personal communication). We therefore mutated these three amino acids to alanines (UPF1 RM, for RNA mutant) and tested the capacity of this mutant to bind RNA using MicroScale Thermophoresis (MST). To this end, we affinity-purified triple-FLAG-tagged UPF1 proteins (UPF1-3F and UPF1 RM-3F) from HEK cells (Figure 3A). The purified recombinant proteins were titrated (from 200 nM to 0.0061 nM) against a constant amount (20 nM) of Cy5-labelled U30 RNA. Binding of wt UPF1 to RNA displayed a sigmoidal dose-response curve from which a *K*_d_ of 25.33 ± 4.43 nM was calculated, whereas UPF1 RM failed to bind the U30 RNA (Figure 3B).

**Figure 3.**
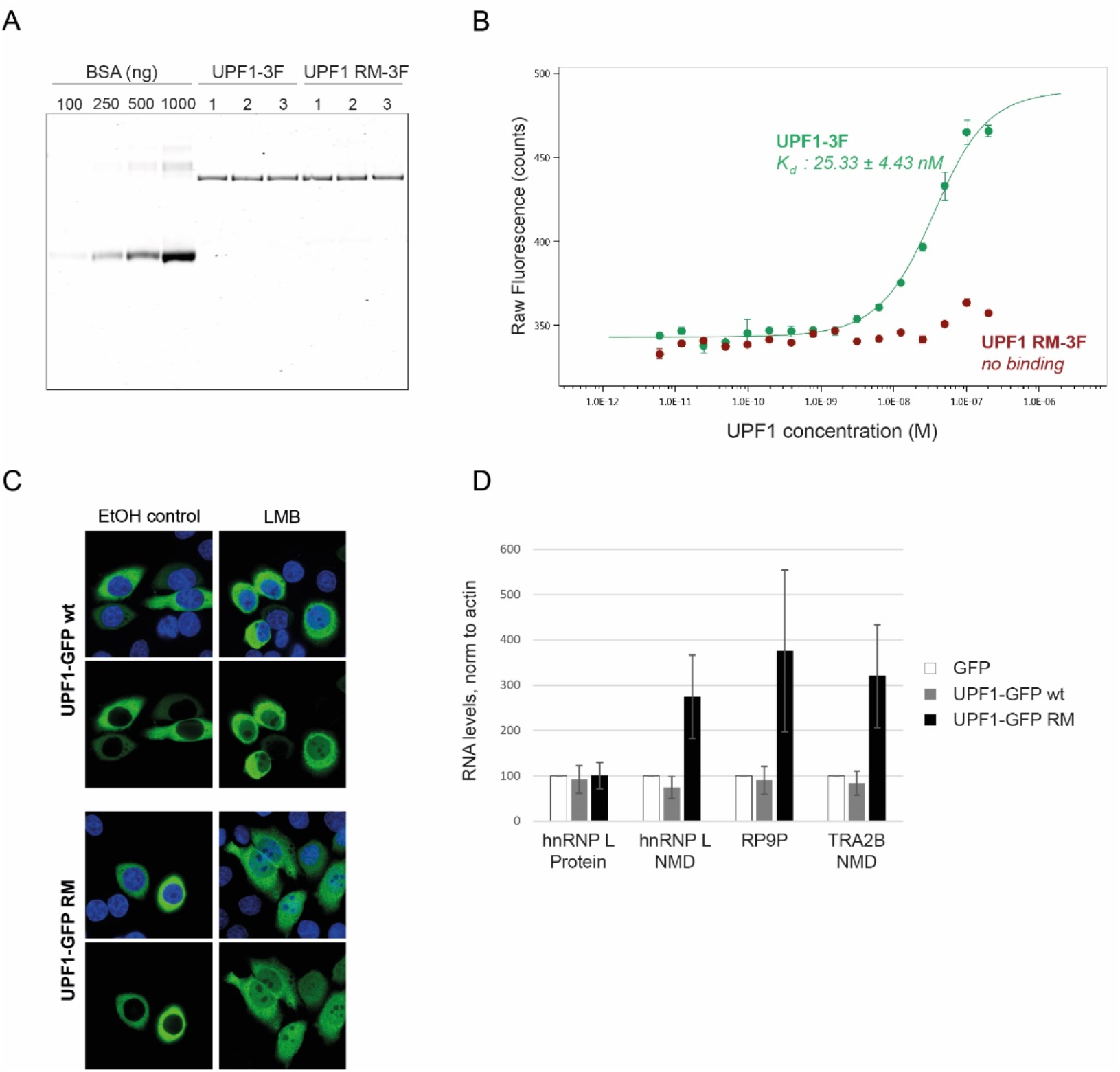
RNA binding is not necessary for shuttling. **A**. UPF1-3F and UPF1 RM-3F (RNA mutant: N524A, K547A and R843A) proteins were overexpressed in HEK cells and affinity purified by the triple Flag tag. The quality of the purification was assessed by SDS-PAGE. 25% of the immunoprecipitated material was loaded in a 4-12% gradient gel and proteins were stained with Imperial Protein Stain. Three independent purifications (1,2,3) are shown. BSA was used to estimate the concentration of the purified proteins. **B**. UPF1-3F (in green) and UPF1 RM-3F (in red) binding to Cy5-labeled U30 RNA (20 nM) was analyzed by MicroScale Thermophoresis. The raw fluorescence counts are plotted against the final concentration of the unlabelled titrated proteins (200–0.0061 nM). The estimated *K*_d_ for UPF1-3F is 25.33 ± 4.43 nM. Averages and standard deviations from two independent measurements are shown. **C**. UPF1-GFP or UPF1-GFP RM were expressed in HeLa cells and GFP signal was monitored by confocal fluorescence microscopy. GFP signal (green channel) and GFP together with DAPI signal (blue channel) are shown in control condition and upon addition of LMB (6 hours). **D**. RNA levels of HeLa cells expressing of GFP, UPF1-GFP wt or UPF1-GFP RM were analyzed by RT-qPCR. Relative mRNA levels of hnRNP L Protein, hnRNP L NMD, RP9P, and TRA2B NMD, normalized to actin mRNA levels, are depicted. Averages and standard deviations arise from three independent experiments.

The localization of this mutant was eventually investigated by fluorescence microscopy analysis and compared to wt UPF1-GFP. Confocal fluorescence microscopy revealed that UPF1-GFP RM was still able to move between cytoplasm and nucleus as judged from the nuclear accumulation upon LMB treatment, indicating that the capacity of UPF1 to shuttle does not depend on its binding to RNA (Figure 3C). Interestingly, this RNA binding mutant of UPF1 had a dominant negative effect on NMD (Figure 3D and below).

### Amino acids L103 and F105 are required for nuclear export of UPF1

To characterize the amino acid residues crucial for nuclear export, the N-terminal region of UPF1 that was previously identified as being required for export (amino acids 55-416) (Mendell *et al*., 2002) was analyzed by the bioinformatic tool ‘LocNES’ (Xu et al., 2015). This algorithm predicted amino acids 89-105 (VDDSVAKTSQLLAELNF), just upstream of the CH domain, to function as a NES. We deleted the nucleotides encoding this 17 amino acids stretch by fusion PCR, generating the plasmid UPF1-GFP Δ89-105. Analysis by confocal microscopy of this UPF1 deletion showed a clear nuclear signal both in the control and LMB condition (Figure 4A). Classical NES (also called leucine-rich NES) are typically 8-15 residues long and characterized by conserved hydrophobic amino acids – often leucine – that are separated by one, two and three other amino acids (e.g. L-X_3_-L-X_2_-L-X-L, where L is a hydrophobic amino acid and X any amino acid) (Kosugi et al., 2008). To inactivate the putative NES, two or four hydrophobic amino acids in the predicted NES of UPF1 were changed to alanine giving raise to the mutant constructs UPF1 LF and UPF1 VLLF, and expression of the proteins in HeLa cells was confirmed by Western blotting (Figure S2A). Analysis of these UPF1-GFP mutants by fluorescence microscopy showed that these amino acid residues are indeed important for the export of UPF1 from the nucleus. Mutations of two (L103A and F105A) or four (V93A, L100A, L103A, and F105A) amino acids led to clear nuclear signals already without LMB, and the signals stayed the same when LMB was added for six hours (Figure 4A).

**Figure 4.**
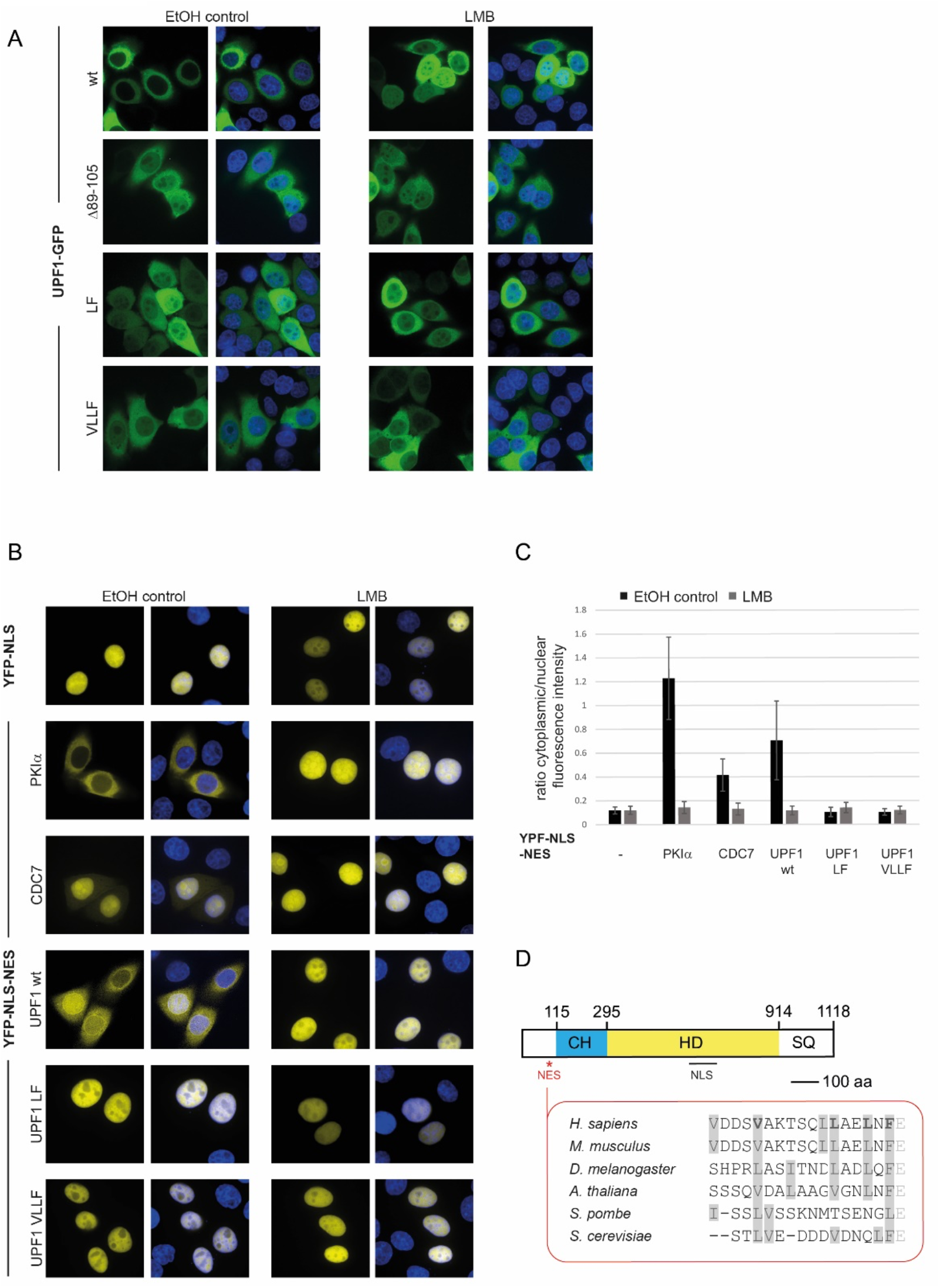
Investigation of UPF1’s NES. **A**. UPF1-GFP wild type or variants (deletion of amino acid 89-105; point mutations V93A, L100A, L103A, F105A and L103A, F105A) were expressed in HeLa cells and analyzed as in Figure 3C. **B**. The localization of YFP was determined by confocal fluorescence microscopy and DAPI was used for staining the nucleus. YFP followed by the NLS of SV40 served as a control (upper panel). Insertion of a NES (of PKIα, CDC7 or UPF1 wt) to YFP-NLS led to a shift of the yellow signal to the cytoplasm, but insertion of a mutated NES (UPF1 LF and UPF1 VLLF) to YFP-NLS or LMB treatment prevented the export of YFP. **C**. The average ratio of mean cytoplasmic to mean nuclear fluorescence intensity are shown with standard deviations. In total 50 cells per condition (from three independent experiments) were analyzed by Cell Profiler software (www.cellprofiler.org). DAPI and phalloidin staining were used for defining nucleus and cytoplasm (see also Figure S2B). **D**. Schematic illustration of UPF1 as in Figure 1A with amino acid sequence of the NES in human, mouse, fly, plant, and yeasts. Hydrophobic amino acids (phenylalanine, leucine, isoleucine, valine) are marked in grey.

To unambiguously confirm (i) that this region of UPF1 is indeed the NES, (ii) that amino acids L103 and F105 are necessary for NES activity, and (iii) that the observed localization is not an effect of misfolded UPF1, a simple but elegant system from the Chook laboratory was adopted (Xu *et al*., 2015). In this reporter system, the strong NLS of SV40 was fused C-terminally to the enhanced yellow fluorescent protein (YFP), resulting in an exclusively nuclear signal in the absence and presence of LMB (Figure 4B). Next, different NESs were added downstream of YFP-NLS to test their ability to re-localize YFP to the cytoplasm. As control for a strong and weak NES sequence, the NES of PKIα and CDC7 were used, which results in a strong or a faint cytoplasmic YFP signal, respectively (Xu *et al*., 2015) (Figures 4B and S2B). To test the putative NES of UPF1 in this assay, the sequence coding for the 17 amino acid residues predicted by LocNES plus the subsequent highly conserved glutamate (VDDSVAKTSQLLAELNFE) was added downstream to YFP-NLS coding sequence and after transient transfection into HeLa cells, the yellow fluorescence was detected by confocal fluorescence microscopy. In the absence of LMB (EtOH control), the putative NES of UPF1 (UPF1 wt) led to a clear shift of the YFP signal towards the cytoplasm, indicating that these amino acids are sufficient to promote nuclear export of the YFP-NLS construct and hence constitute a functional NES (Figure 4B). Quantitative fluorescence intensity analysis of 50 cells per condition using the Cell Profiler software (Carpenter et al., 2006) revealed that the UPF1 NES caused a 6-fold increase in the fluorescence intensity ratio between cytoplasm and nucleus compared to the YFP-NLS control, which indicates that the strength of the UPF1 NES is somewhere in between the strong PKIα NES (10-fold increase) and the weaker CDC7 NES (3.5-fold increase) (Figure 4C). Importantly, addition of LMB prevented the export of the YFP protein with each of the three NES, confirming that they engage the CRM1-dependent export pathway (Figure 4B-C).

When the two mutated versions of the UPF1 NES (UPF1 LF: VDDSVAKTSQLLAE**A**N**A**E and UPF1 VLLF: VDDS**A**AKTSQL**A**AE**A**N**A**E) were tested in this assay, no export of YFP to the cytoplasm was observed, confirming that amino acids L103 and F105 are necessary for UPF1’s NES function (Figures 4B-C and S2B). Collectively, we have precisely defined the NES in UPF1 and identified two crucial amino acids (L103 and F105) that when mutated inactivate this NES. Notably, the UPF1 NES sequence is highly conserved among mammals, *D. melanogaster*, and *A. thaliana* but not so clearly defined in *S. pombe* (Figure 4D).

### The LF mutant is functional in NMD and interacts with known NMD factors

In the next series of experiments, we aimed to characterize if a functional NES is critical for the activity of UPF1. To this end, we used the above-mentioned NES mutants and analyzed whether the interaction of UPF1 to other proteins is disturbed by inactivation of the NES. Immunoprecipitation experiments using GFP nanobodies in HeLa cells expressing either GFP alone, UPF1-GFP wt, UPF1-GFP LF or UPF1-GFP VLLF revealed no difference in the interaction pattern between the wt and the mutated UPF1 proteins (Figure S2C). When we examined the immunoprecipitated material by Western blotting, interactions with the two NMD factors UPF2 and SMG6 were confirmed for wt UPF1 and for the NES mutants LF and VLLF (Figure 5A). Specificity of the immunoprecipitation was controlled in HeLa cells expressing GFP alone and the absent interaction to GAPDH for all pulldowns (Figure 5A).

**Figure 5.**
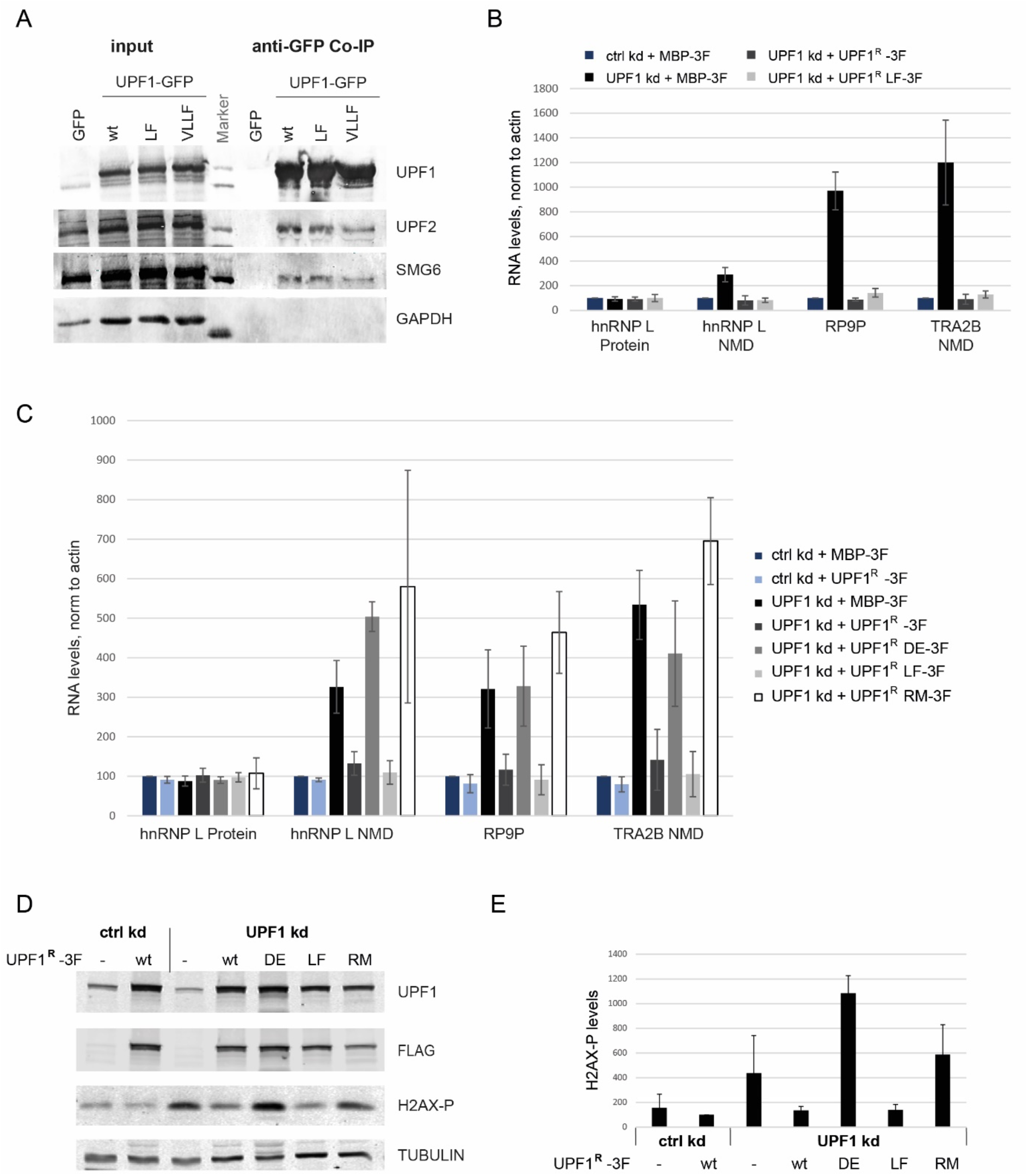
UPF1 LF mutant is functional in NMD. **A**. Western Blotting to analyze co-immunoprecipitation experiments performed with the help of GFP nanobodies in HeLa cells expressing GFP alone, UPF1-GFP wt, UPF1-GFP LF mutant or UPF-GFP VLLF mutant. 2 % of starting material (input) and 60% of the immunoprecipitated material was loaded on an SDS-PAGE. The membrane was probed against UPF1, UPF2, SMG6, and GAPDH as a control. **B**. HeLa cells were transfected with pKK plasmids to simultaneously knockdown UPF1 (UPF1 kd) and over express RNAi resistant UPF1 wt or LF mutant. Knockdown against luciferase (ctrl kd) and expression of maltose binding protein (MBP) served as control. The cells were selected with puromycin for 48 hours and harvested 96 hours after transfection. Relative mRNA levels of hnRNP L Protein, hnRNP L NMD, RP9P, and TRA2B NMD, normalized to actin mRNA levels, were determined by RT-qPCR. Averages and standard deviations from three independent experiments are shown. **C**. HEK cells were induced for 96 h with Dox to express simultaneously siRNAs against UPF1 (UPF1 kd) or against luciferase (ctrl kd) and RNAi resistant UPF1 versions (wt, DE, LF or RM variants). As control, MBP was expressed instead of UPF1^R^. Relative mRNA levels, normalized to actin mRNA levels, were measured by RT-qPCR in three independent experiments as in B. **D**. H2AX phosphorylation was assessed by immunoblotting of HEK cells harvested 96 hours after addition of Dox (as in C). UPF1 expression (endogenous and Flag-tagged) was controlled by probing the membrane with anti-UPF1 and anti-FLAG antibody. TUBULIN served as loading control. **E**. Phosphorylated H2AX levels on Western Blots were quantified by using Image Studio software and normalized to TUBULIN. Averages and standard deviations of three independent experiments are depicted.

Finding that the NES-impaired mutant still interacts with important NMD factors and that the mutations are not part of the CH or helicase domain led us to hypothesize that the LF mutant might still function in NMD. To test this, we used a plasmid with a bidirectional promoter that allows simultaneous knockdown (kd) of the endogenous UPF1 protein and expression of an RNAi resistant UPF1 protein, either wt or mutant versions thereof. We expressed the plasmids transiently in HeLa cells (Figure 5B and S3A) or stably integrated them into HEK293-T Rex Flp-in cells (Figure 5C-E and S3B-D). All UPF1 constructs were tagged at the C-terminus with a triple FLAG peptide for affinity purification (Figure 3A), immunofluorescence (Figure S3A-B) and Western blot analysis (Figure 5D). Immunofluorescence experiments clearly revealed that, while a fraction of the UPF1 LF mutant localized to the nucleus, most of it is still found in the cytoplasm where the protein might be needed for the translation-dependent NMD pathway (Figure S3A-B).

In the HEK293 cells, the bidirectional promoter was induced by doxycycline (Dox) and cells were analyzed at different time points to find the best knockdown and rescue conditions. 96 hours after Dox addition, endogenous UPF1 protein was robustly reduced, the stabilization of NMD transcripts in the knockdown condition was the highest for two out of three tested transcripts, and NMD activity could be efficiently rescued by the expression of a FLAG tagged UPF1 wt version (UPF1^R^-3F) (Figure S3C-D). Analysis of the different HEK cell lines by RT-qPCR 96 hours after Dox induction showed increased RNA levels for endogenous NMD-sensitive transcripts (hnRNP L NMD, RP9P, and TRA2B NMD) when the endogenous UPF1 was depleted and an unrelated control protein (maltose-binding protein, MBP-3F) expressed (Figure 5C). The observed stabilization of transcripts is specific for NMD targets, since transcripts that are not targeted by NMD like the hnRNP L protein coding splice isoform (hnRNP L Protein) or actin did not react to the UPF1 kd. As expected, expression of RNAi-resistant wt UPF1 was able to rescue the NMD phenotype and the stabilization was reversed to control levels (UPF1 kd + UPF1R-3F; Figures 5C and S3D). The helicase-deficient DE mutant served as a control, since the helicase activity of UPF1 is required for NMD and therefore, as expected, NMD activity was not restored when overexpressing this mutant (UPF1 kd + UPF1^R^ DE-3F; Figure 5C). Interestingly, the NES-inactivating LF mutant was fully functional in NMD and restored NMD activity in cells with a kd of endogenous UPF1 (UPF1 kd + UPF1^R^ LF-3F; Figure 5C). The functionality of the LF mutant in NMD was corroborated when transiently expressed in HeLa cells (Figure 5B). The new RNA binding mutant characterized in Figure 3 was also not able to rescue the UPF1 kd phenotype and RNA levels of the NMD sensitive transcripts are similar to the ones in the DE mutant (UPF1 kd + UPF1^R^ RM-3F; Figure 5C). It should be noted that overexpression of this mutant was detrimental for cells and, without knocking down the endogenous UPF1, the RNA binding mutant acts as dominant negative mutant (Figure 3D). Notably, one of the three amino acids that were changed was already shown in 1998 to have dominant-negative effects on NMD (Sun et al., 1998).

Lastly, the function of the LF mutant in the phosphorylation of H2AX was investigated. Nuclear functions of UPF1 are poorly understood and it is not clearly proven whether H2AX phosphorylation is a direct effect of UPF1 in the nucleus or an indirect effect of for instance impaired NMD occurring in the cytoplasm. As reported before, depletion of UPF1 resulted in an increase of H2AX phosphorylation and this increase can be rescued with the wt version of UPF1 (Azzalin and Lingner, 2006) (Figure 5D-E). The NES mutated LF version acts very similar to UPF1 wt leading to a reduced H2AX phosphorylation, similar to control levels, and is thus functional in this process. The DE and the RNA binding mutant were not able to reverse the H2AX increase and are therefore not only impaired in the NMD pathway, but also in this nuclear outcome (Figure 5C-D).

## Discussion

Although there is ample evidence for UPF1 to be a shuttling protein with only a small fraction of the protein residing inside the nucleus at steady state, the NLS and NES of UPF1 have so far not been precisely mapped. In this study, we first showed that the intracellular distribution is indistinguishable between the two UPF1 isoforms 1 and 2, which differ by an 11 amino acid loop in the helicase domain, and then confirmed a previously reported 100 amino acids stretch in the helicase domain to be necessary for nuclear import of UPF1 (Mendell *et al*., 2002). Further delineation of this NLS, however, was not possible since a smaller deletion of that region (Δ596-606) or mutations therein (A1-A5) did not lead to any changes in localization of the protein compared to the wild type situation. By contrast, the LocNES algorithm (Xu *et al*., 2015) successfully predicted the NES to reside between amino acids 89 and 105, in the N-terminal part of UPF1 right upstream of the CH domain (Figure 3D). Changing the conserved amino acids L103 and F105 to alanine abolished the activity of this NES and led to UPF1 accumulation in the nucleus.

We noticed that based on the biochemical fractionation between cytoplasm and nucleus, a larger fraction of the total cellular UPF1 appeared to be present in the nucleus at steady state than when compared to the amount of nuclear UPF1 detected on the fluorescence microscopy images (Figure 1B and C). We attribute this to the fact that nuclear fractions, in addition to intranuclear material, tend to also contain proteins associated with the outer nuclear membrane and it was recently shown that a fraction of the cellular UPF1 associates with the endoplasmic reticulum (ER) (Longman et al., 2020), which is seamlessly connected to the outer nuclear membrane. Consistent with this view, the microscopic images often show a ring-like structure around the nucleus, indicating a high concentration of UPF1 around it (Figures 1 and 2). Notably, the two UPF1 isoforms 1 and 2, which differ by 11 amino acids in a regulatory loop of the helicase domain, showed the same subcellular localization (Figure 1C), arguing against isoform-specific nuclear and cytoplasmic functions of UPF1. Since the NLS and the NES that we defined in this study do not overlap with the alternatively spliced part that gives raise to the two UPF1 isoforms, this observation was expected.

The experiments with the export inhibitor LMB showed that UPF1 is imported to the nucleus at a rather slow rate and even 6 hours after LMB addition not all UPF1 localized to the nucleus, indicating that the ill-defined NLS of UPF1 must be considerably weaker than a classical NLS. Perhaps, the NLS of UPF1 is not always accessible for the import receptor and a sub-population of UPF1 might not be imported to the nucleus because its NLS might be masked by other interacting proteins or by an alternative conformation of UPF1 itself. Despite all efforts, we were not able to pinpoint the NLS of UPF1 to specific amino acids or to a shorter amino acid stretch than previously reported (Mendell *et al*., 2002). In addition to the common importin α-dependent import, there are cases reported where importin β directly binds to non-canonical NLSs that are more diverse and less defined (Lee et al., 2006). Many of these non-canonical NLSs directly bound by importin β1 are also rich in arginine and therefore mutating positively charged amino acids in the helicase region 607-697, as done in Figure 2B, would most likely also disturb nuclear import mediated by importin β1. Non-canonical NLSs that are recognized by importin β2 have often an overall basic character plus a relatively conserved proline-tyrosine (PY) dipeptide (Lee *et al*., 2006). However, no such a PY-NLS could be identified in UPF1, leading to the speculation that UPF1 might hitchhike to the nucleus by binding to another protein.

Interestingly, helicase deficient D636A/E637A UPF1 stayed cytoplasmic even when cells were treated with LMB, corroborating the results obtained with *Drosophila* cells (Singh *et al*., 2019). Since UPF1 helicase activity is required for its dissociation from RNA, we hypothesized that the impaired helicase activity leads to formation of cytoplasmic protein-RNA aggregates that would trap UPF1 in the cytoplasm (Franks *et al*., 2010). In fact, we showed here that UPF1’s subcellular localization is not influenced by its RNA binding capacity, as the triple mutant N524A/K547A/R843A, which no longer binds RNA, did not alter the shuttling behaviour. The RNA binding mutant still accumulated in the nucleus upon LMB treatment, indicating that the import of UPF1 does not require its capacity to bind RNA. Furthermore, its cytoplasmic localization at steady state, indistinguishable from wild type UPF1, indicates that association with RNA is not required for the export of UPF1. Supporting this conclusion is also the observation that UPF1 export is inhibited by LMB, implying that the export of UPF1 is mediated by CRM1 rather than by the main mRNA export factor NXF1 (Okamura et al., 2015). Collectively, the results of these mutants showed that the helicase activity and the RNA binding capacities are indispensable for NMD but do not affect the subcellular distribution of UPF1.

In this study we mapped the NES of UPF1 to amino acids 89-105 and showed that L103 and F105 are crucial for its activity, since replacement of these two amino acids with alanine led to a clear redistribution of UPF1 signal towards the nucleus. Notably, this region is highly conserved among metazoans but slightly less in the yeasts *S. pombe* and *S. cerevisiae* (Figure 4D), suggesting that the shuttling behaviour of UPF1 might be restricted to metazoans or that in yeast, another NES in another part of the protein has evolved.

Having identified the NES of UPF1 allowed us to increase UPF1 abundance inside the nucleus, which we assumed might affect both nuclear and cytoplasmic functions of UPF1. We found that the NES-deficient UPF1 could rescue NMD to the same extent as wild type UPF1 in cells depleted for endogenous UPF1, which is not surprising given that still a large amount of the NES mutated protein localized to the cytoplasm. With regards to a reported nuclear function of UPF1, we followed the approach taken by Azzalin and Lingner (Azzalin and Lingner, 2006) who showed that knocking down UPF1 led to increased H2AX phosphorylation, suggesting a direct or indirect role for UPF1 in DNA damage response. When expressing the different UPF1 mutants in cells depleted of endogenous UPF1, the NES-defective mutant was as good as wild type UPF1 in rescuing H2AX phosphorylation, whereas the helicase and the RNA binding mutants were unable to rescue. Curiously, the result of the H2AX phosphorylation experiment mirrored the ability of the tested UPF1 mutants to rescue NMD (Figure 5), which could indicate either that the DNA damage response and NMD are somehow mechanistically linked, or that they are independent of each other but coincidentally depend both on the same properties of UPF1 (helicase activity and RNA binding). Further investigations are needed to clarify whether UPF1’s reported roles in nuclear processes are direct or indirect.

## Material and Methods

### Plasmids

Oligonucleotides used are listed in Table S1. To generate <u>UPF1-GFP iso2</u> a *Xba*I restriction site was introduced in pcDNA3-NG-UPF1-WT-FLAG (Rufener and Muhlemann, 2013) downstream of FLAG tag by site-directed mutagenesis. GFP was amplified by PCR from pcDNA3-MS2-EGFP (McNally et al., 2006) using oligonucleotides introducing *Not*I and *Xba*I sites. The GFP amplicon was then ligated into *No*tI and *Xba*I cut plasmid pcDNA3-NG-UPF1-WT-FLAG. Annealed oligonucleotides encoding a linker of 16 glycine residues (see sequence in Table S1) were inserted into the vector at the *Not*I site to receive the plasmid encoding the fusion protein UPF1-GFP with a flexible linker in-between. For obtaining <u>UPF1-GFP iso1</u>, a fragment of UPF1 (from *Hind*III to *Kpn*I) was replaced in UPF1-GFP iso2 with the corresponding iso1 UPF1 fragment. <u>UPF1-GFP NLS (SV40)</u> and <u>UPF1-GFP NES (snurportin-1)</u> were generated by digesting UPF1-GFP iso2 plasmid with *BsrG*I and *Xba*I (both sites located at the very end of GFP) and purifying it by Wizard® SV Gel and PCR Clean-Up System (Promega). Annealed oligonucleotides encoding the NLS of SV40 or NES of snurportin-1 with a stop codon and *BsrG*I and *Xba*I overhangs at the end are ligated into the linearised UPF1-GFP iso2 vector (oligonucleotides listed in Table S1). Fusion PCR was performed to generate <u>UPF1-GFP Δ89-105, Δ596-697, and Δ596-606</u> variants using the oligonucleotides listed in Table S1. <u>UPF1-GFP DE</u> was cloned by exchanging a part of UPF1-GFP iso2 (from *Kpn*I to *Sbf*I) with the same part of pcDNA3-NG-UPF1-WT-FLAG DE636AA containing the two desired changes. Point mutations for the <u>UPF1-GFP RM</u> were introduced individually by site-directed mutagenesis PCR on cloning plasmids containing parts of UPF1 and then mutated UPF1 sequences were cloned into the final plasmid by restriction sites (oligonucleotides for point mutations are listed in Table S1). Point mutations in UPF1-GFP to generate <u>UPF1-GFP A1-A6</u> were introduced by synthesising a part of UPF1 with the desired mutations (General Biosystems, Inc) and these fragments were then used to replace the original sequence. *Kpn*I and *Sbf*I were used as restrictions sites for exchanging part of the UPF1 sequence. <u>UPF1-GFP LF and UPF1-GFP VLLF</u> was generated by synthesis a piece of UPF1 (from *Hind*III to *Kpn*I) with the desired point mutations (General Biosystems, Inc). The *Hind*III-*Kpn*I fragment was then introduced into the plasmid encoding UPF1-GFP iso2 digested by *Hind*III and *Kpn*I. The <u>YFP-NLS plasmids</u> (without NES and with NES of PKIα and CDC7) were a kind gift from Yuh Min Chook’s laboratory (Xu *et al*., 2015). To clone UPF1’s wild-type or mutant NES at the C-terminus of YFP-NLS, YFP-NLS plasmid was digested by *Bam*HI and *Hind*III and purified by Wizard® SV Gel and PCR Clean-Up System (Promega). Annealed oligonucleotides encoding UPF1’s NES with *Hind*III and *Bam*HI overhangs at the end (UPF1-NES wt, UPF1-NES LF or UPF1-NES VLLF, listed in Table S1) are then ligated into the linearized vector.

<u>pKK plasmids</u> expressing from bidirectional, Tet responsive CMV promoter an RNAi cassette (luciferase as control or UPF1 cassette) on one side and the protein of interest (triple FLAG tagged MBP or UPF1) on the other side were designed and cloned following (Szczesny et al., 2018). Each RNAi cassette contains three siRNAs which contain self-complementary regions and a loop-forming sequence from murine miRNA 155. For designing the RNAi cassette of (firefly) luciferase (*P. pyralis*, NCBI accession number M15077.1) LOCK-iT™ RNAi Designer from Thermo Fisher Scientific (https://bit.ly/3qcESry) was used and for UPF1 RNAi cassette the already well-established siRNA targets were chosen (Paillusson et al., 2005). The RNAi cassettes were synthesised (General Biosystems, Inc) and inserted into the pKK vector with the help of *Apa*I and *Mfe*I restriction sites. C-terminally triple FLAG tagged MBP was inserted into pKK plasmids using InFusion cloning (Takara). To introduce C-terminally triple FLAG tagged UPF1, the pKK vector was partially digested with *Not*I and *Hind*III (since two *Not*I sites are present) and the desired vector backbone was purified via agarose gel. RNAi resistant UPF1 was obtained from cutting pcDNA3-NG-UPF1-WT-FLAG (Rufener and Muhlemann, 2013) with *Not*I and *Hind*III and cloned into the linearized vector backbone. To obtain pKK plasmids with variant UPF1 (pKK UPF1 kd + UPF1^R^ DE, + UPF1^R^ LF, + UPF1^R^ RM) pieces of the wt UPF1 were replaced by the mutant ones using *Hind*III and *Pml*I restriction sites. For transient transfection of pKK plasmid in HeLa cells, the hygromycin resistance was replaced by puromycin resistance cassette.

For all PCR reactions, high fidelity polymerases were used (Kapa Hifi Hot Start Ready mix (Roche) or CloneAmp HiFi PCR Premix (Takara)) and sequences of plasmids were confirmed by Sanger sequencing (Microsynth, Switzerland).

### Cell culture and transfection

HeLa cells and HEK293 Flp-In™ T-Rex cells (Thermo Fisher Scientific) were cultured in Dulbecco’s Modified Eagle Medium (DMEM) supplemented with 10% fetal bovine serum (FBS), 100 U/ml penicillin and 100 *μ*g/ml streptomycin (P/S) at 37 °C under 5% CO_2_.

To stably introduce pKK plasmids into HEK293 Flp-In™ T-Rex cells, 3 x 10^6^ cells were seeded in a 10-cm plate using DMEM plus tetracycline-free FBS. Next day, 2 *μ*g of the pKK plasmid and 18 *μ*g of pOG44 Flp-Recombinase expression vector (Thermo Fisher Scientific) were transfected with the help of lipofectamine 2000 (Thermo Fisher Scientific). 24 hours after transfection, cells were transferred into T150 flasks (DMEM + tetracycline-free FBS) and 3 hours later the antibiotics are added (P/S, 100 *μ*g/ml hygromycin B, 10 *μ*g/ml blasticidine). Cells were kept under selection for about two weeks. Transcription of the plasmids was induced by the addition of 1 *μ*g/ml doxycycline (Dox).

For transient transfection in HeLa cells, 2 x 10^5^ cells were seeded in a 6-well plate and 500 ng UPF1-GFP plasmids (wt or variants of it, for Figure 1C, 1D, 2, 3C-D, 4A) or 500 ng YFP-NLS plasmids (for Figure 4B) were transfected the next day using Dogtor transfection reagent (OZ Biosciences). 24 hours later transfection efficiencies were evaluated by using fluorescence microscopy and 40’000 cells were seeded on 8-well chamber slides for fluorescence microscopy analysis the next day. Samples for Western Blotting and RNA analysis were also harvested 48 hours after transfection.

For transient rescue experiments in HeLa cells (Figure 5B, S3A), 1000 ng of pKK plasmids were transfected with Dogtor transfection reagent in wells of 6-well plates. The next day cells were transferred to T25 flasks and selected for 48 hours with 1.5 *μ*g/ml puromycin. Before analysing the experiment by immunofluorescence and RT-qPCR, cells were cultivated one additional day without puromycin.

### LMB treatment, detection of GFP/YFP, and fluorescence microscopy analysis

Cells in 8-well chamber slides were incubated for 6 hours with 50 nM LMB (Cell Signaling Technology) or with ethanol as control. After washing cells with PBS twice, cells were fixed with 4 % paraformaldehyde for 25 minutes at room temperature. Next, cells were washed three times with TBS (20 mM Tris-HCl, pH 7.5, 150 mM NaCl) and incubated with TBS +/+ (TBS plus 0.5 % Triton X-100 and 6 % FBS) containing DAPI and 594-Phalloidin (Hypermol, for Figure 4B and S2B) for 1 hour at 37 °C. Finally, cells were washed three times with TBS and Vectashield mounting medium (Vector Laboratories) was added on slides to mount the coverslips. The slides were analyzed by epifluorescence microscope (Leica Microsystems, DMI6000 B) or confocal spinning disc microscopy (Nikon TI2 with Crest X-Light V2). CellProfiler software was used for quantifications of images [https://cellprofiler.org/ (Carpenter et al., 2006)].

### Biochemical fractionations of HeLa cells

HeLa cells were counted to keep 2 x 10^5^ cells for *whole cells* input sample. 1 x 10^7^ cells were spun down (500 x g, 5 min), the pellet was resuspended in cold buffer 1 (50 mM HEPES, pH 7.6, 140 mM NaCl, 1 mM EDTA, 10 % glycerol, 0.5 % NP-40, 0.25 % Triton X-100 and protease inhibitors) and incubated for 10 minutes on a rotating wheel in the cold room. Cells were centrifuged as before, and the supernatant was kept as *cytoplasmic fraction*. This first step was repeated twice to ensure total cell lysis. Next, the pellet was resuspended in cold buffer 2 (10 mM Tris, pH 8, 200 mM NaCl, 1 mM EDTA, 0.5 mM EGTA and protease inhibitors) and incubated again 10 minutes. After centrifugation, the nuclei were solubilized in cold buffer 3 (10 mM Tris, pH 8, 1 mM EDTA, 0.5 mM EGTA, 0.1 % SDS, 0.1 % deoxycholate, 1 % Triton X-100 and protease inhibitors) and sonicated briefly by tip sonication (Vibra-Cell™ 75186, 3 x 10 seconds, 45 % amplitude). Finally, samples were centrifuged (16’000 x g, 30 minutes, 4° C) and the supernatant was kept as *nuclear fraction*. For Western Blot analysis, proteins of the cytoplasmic and nuclear fraction (corresponding to 2 x 10^5^ cells) were precipitated by ice-cold acetone.

### Co-immunoprecipitation

For investigating interaction partners of UPF1, 12 *μ*g of UPF1-GFP plasmid (wt or variants of it) was transfected with the transfection reagent Dogtor (OZ Biosciences) into HeLa cells seeded the day before in a 15-cm plate. 48 hours after transfection, cells were harvested, lysed in cold lysis buffer (50 mM HEPES, pH 7.3, 150 mM NaCl, 0.5 % Triton X-100 and protease inhibitors) and sonicated briefly by tip sonication (Vibra-Cell™ 75186, 3 x 10 seconds, 45 % amplitude). After centrifugation (16’000 x g, 10 min, 4° C), the supernatant was incubated with magnetic GFP-TRAP beads (Chromotek) on a rotating wheel for one hour in the cold room. After washing the beads twice on a magnet with lysis buffer, the samples were incubated with 100 *μ*g RNase A for 10 minutes at 25 °C. The samples were washed once again, and finally LDS-loading buffer was added to the beads. The samples were heated up and loaded on an SDS-PAGE.

### SDS-PAGE, Silver Staining and Western Blotting

was performed as described in (Contu et al., 2021). The Western blots were visualised with the Odyssey System (LICOR) and signals were quantified by Image Studio Software. For mass spectrometry, the prominent band was cut out, destained with Na_2_S_2_O_3_ and K_3_Fe(CN)_6_ and then submitted to the Proteomics Mass Spectrometry Core Facility (PMSCF) of the University of Bern for tryptic in-gel digestion and mass spectrometry analysis.

### Antibodies for Western Blotting

Primary antibodies were diluted in TBS containing 5% BSA or 5% milk and incubated overnight in the cold room. The following antibodies and dilutions were used: goat anti-UPF1 (1:10’000; Bethyl A300-038A;), mouse anti-GAPDH (1:10’000; Santa Cruz Biotechnology sc-47724), rabbit anti-HISTONE H3 (1:10’000; Abcam ab1791), mouse anti-TUBULIN (1:10’000; Sigma-Aldrich T9028), mouse anti-phospho-HISTONE H2A.X (Ser139) (1:1’000; Merck 05-636), mouse anti-FLAG M2 (1:2’000; Sigma-Aldrich F1803), rabbit anti-UPF2 (1:1’000; Bethyl A303-929A) rabbit anti-SMG6 (1:500; Abcam ab87539).

### Immunofluorescence

2 – 4 x 10^4^ HeLa or HEK 293T cells were seeded one day before analysis on an 8-well chamber slides. Cells were fixed with 4 % formaldehyde in PBS for 20 minutes. After washing three times with PBS, cells were permeabilized for one hour with 0.5 % Triton X-100 and 6 % FBS in PBS and further incubated for another hour with mouse anti-FLAG M2 (1:500; Sigma-Aldrich F1803) in PBS with 0.1 % Triton X-100 and 6 % FBS (room temperature). Cells were washed three times with PBS and incubated for one hour with secondary antibody (Alexa Fluor 594, chicken anti-mouse 1:1’000, Invitrogen) in PBS with 0.1 % Triton X-100 and 6 % FBS. Finally, cells were washed as above, stained with DAPI for 10 minutes, washed again and cover slip mounted by using Vectashield mounting medium (Vector Laboratories). The cells were analyzed by confocal spinning disc microscopy (Nikon TI2 with Crest X-Light V2).

### RNA analysis by RT-qPCR

Total RNA was extracted using TRI-mix and precipitated with isopropanol as described in (Nicholson et al., 2012). RNA concentration was measured by Nanodrop and 1 to 3 *μ*g of RNA was reverse transcribed with AffinityScript Multi-Temp RT (Agilent) according to the manufacturer’s protocol. The cDNA was measured by qPCR using Brilliant III Ultra-Fast SYBR Green qPCR mix (Agilent) in the Rotor-Gene Q (Qiagen). The purity of RNA samples was controlled by running RT controls in each experiment. Oligonucleotides for qPCRs are listed in Table S1.

### RNA-protein interaction studies using MicroScale Thermophoresis

#### Purification of recombinant proteins

UPF1^R^-3F and UPF1^R^ RM-3F proteins were expressed in HEK293 Flp-In™ T-Rex cells stably transfected with the respective pKK plasmids and purified by immunoprecipitation. Expression of the recombinant proteins was induced by addition of 1 µg/ml Dox for 48 hours. Cell pellets consisting of 5 x 10^7^ cells were lysed in 500 µl of lysis buffer (500 mM NaCl, 0.5% Triton X-100 and 50 mM Hepes pH 7.4, supplemented with protease inhibitors) and sonicated briefly by tip sonication (Vibra-Cell™ 75186, 3 x 5 sec, 45 % amplitude). After centrifugation (16’000 x g, 10 min, 4° C), the supernatant was incubated with 10 µl of Dynabeads® M-270 Epoxy beads (Invitrogen) coupled with anti-FLAG M2 antibodies on a rotating wheel for two hours at 4 °C. After washing the beads three times with 1 ml of lysis buffer, the proteins were eluted by addition of 20 µl of elution buffer (1 mg/ml 3xFLAG peptide in 1x HBS-EP+ buffer (Cytiva, BR100826)) and incubation at room temperature for 15 min. The purity of the eluted proteins was assessed by SDS-PAGE and Imperial Protein Stain (Thermo Scientific) staining. A BSA standard curve run in the same gel was used to estimate the concentration of the purified proteins.

#### MicroScale Thermophoresis

The RNA-binding activity of UPF1^R^-3F and UPF1^R^ RM-3F was assessed using MicroScale Thermophoresis as described in (Nicholson *et al*., 2014). The 5’-labeled Cy5 RNA U30 oligonucleotide (Microsynth) was diluted to 40 nM in 1x HBS-EP+ buffer. The unlabeled recombinant proteins were serially diluted from a concentration of 200 nM down to 0.0061 nM in the presence of a constant amount of labelled RNA (20 nM). The RNA-protein mixtures were analyzed in premium capillaries using the Monolith NT.115T MST (NanoTemper Technologies). Measurements were carried out at 25 °C, with the following settings: LED power 20%, MST power 40%, laser on time 30 sec, laser off time 5 sec. The NanoTemper Technologies Analysis software, MO.Affinity Analysis v2.3, was used to determine the corresponding *K*_*d*_ using the K_d_ fit model.

## Acknowledgments

We are grateful to Sutapa Chakrabarti (FU Berlin) for pointing out UPF1 residues important for RNA binding, to Lukas Gurzeler and Evan Karousis for helpful discussions throughout the entire project, to Michal Domanski for advice and help in purifying UPF1 and to Julia Schnider, Gianna Fetz and Yuh Min Chook for the UPF1-GFP, pKK puro and pEYFP plasmids, respectively. This work has been supported by the National Center of Competence in Research (NCCR) on RNA & Disease funded by the Swiss National Science Foundation (SNSF), by SNSF grants 31003A-162986 and 310030B-182831 to O.M., and by the canton of Bern (University intramural funding to O.M.).

## Supplementary Information

**Figure S1.**
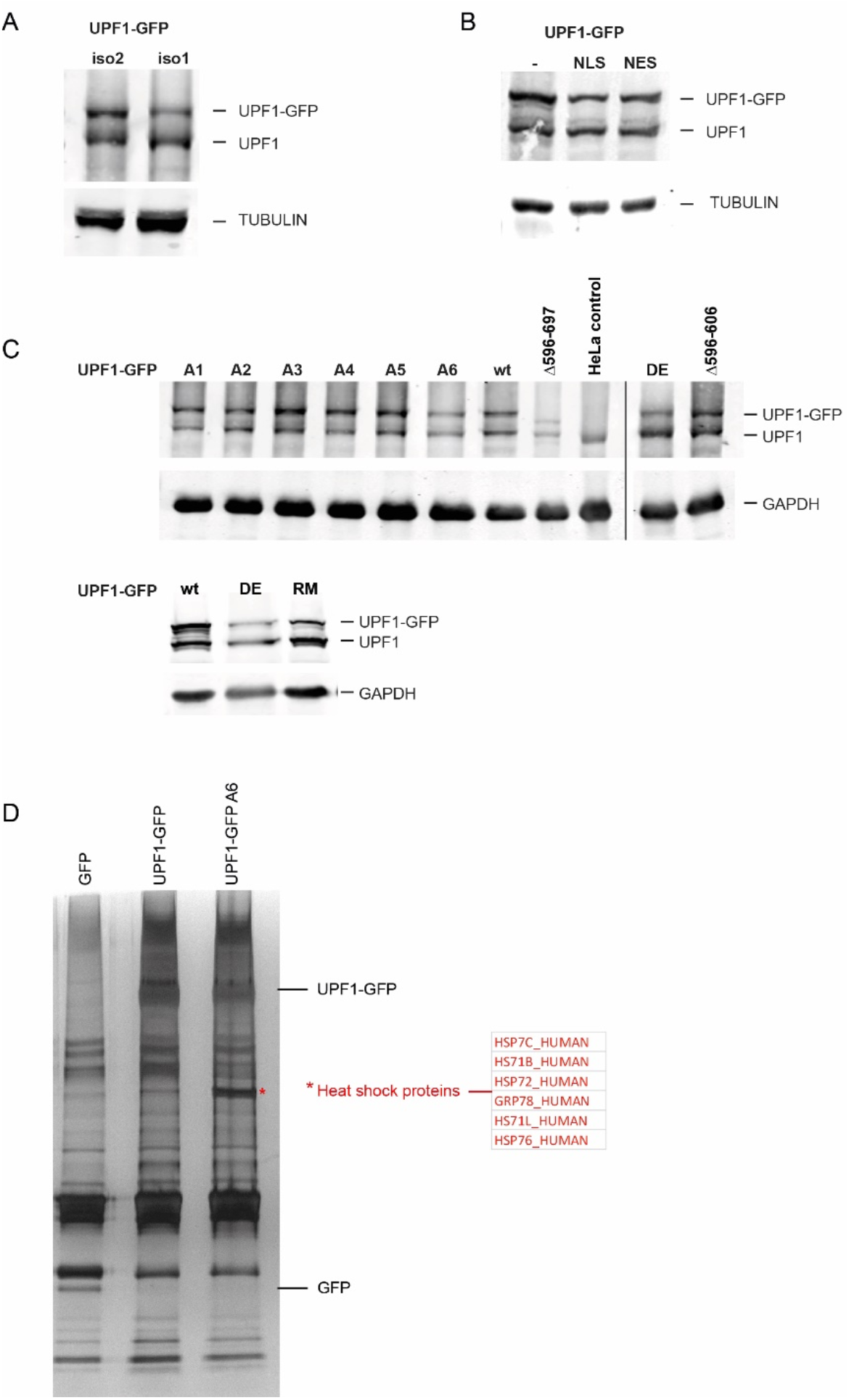
**A**. Expression of UPF1-GFP isoform 1 or isoform 2 in HeLa cells were controlled by probing a Western blot with anti-UPF1 antibody. Tubulin served as loading control. **B**. Immunoblotting to confirm the expression of UPF1-GFP NLS (SV40) and UPF1-GFP NES (snurportin-1) used in Fig. 1D. **C**. The expression of different UPF1-GFP constructs investigated in Figs. 2 and 3 were monitored by Western blotting as in A. GAPDH was used as loading control. **D**. Co-immunoprecipitation experiments of HeLa cells expressing GFP, UPF1-GFP or UPF1-GFP A6 were analyzed by SDS-PAGE and silver staining. Mass spectrometry of the prominent band (marked by red asterisk) appearing at 70 kDa in UPF1-GFP A6 revealed the presence of several heat shock proteins (compared to the same area in UPF1-GFP).

**Figure S2.**
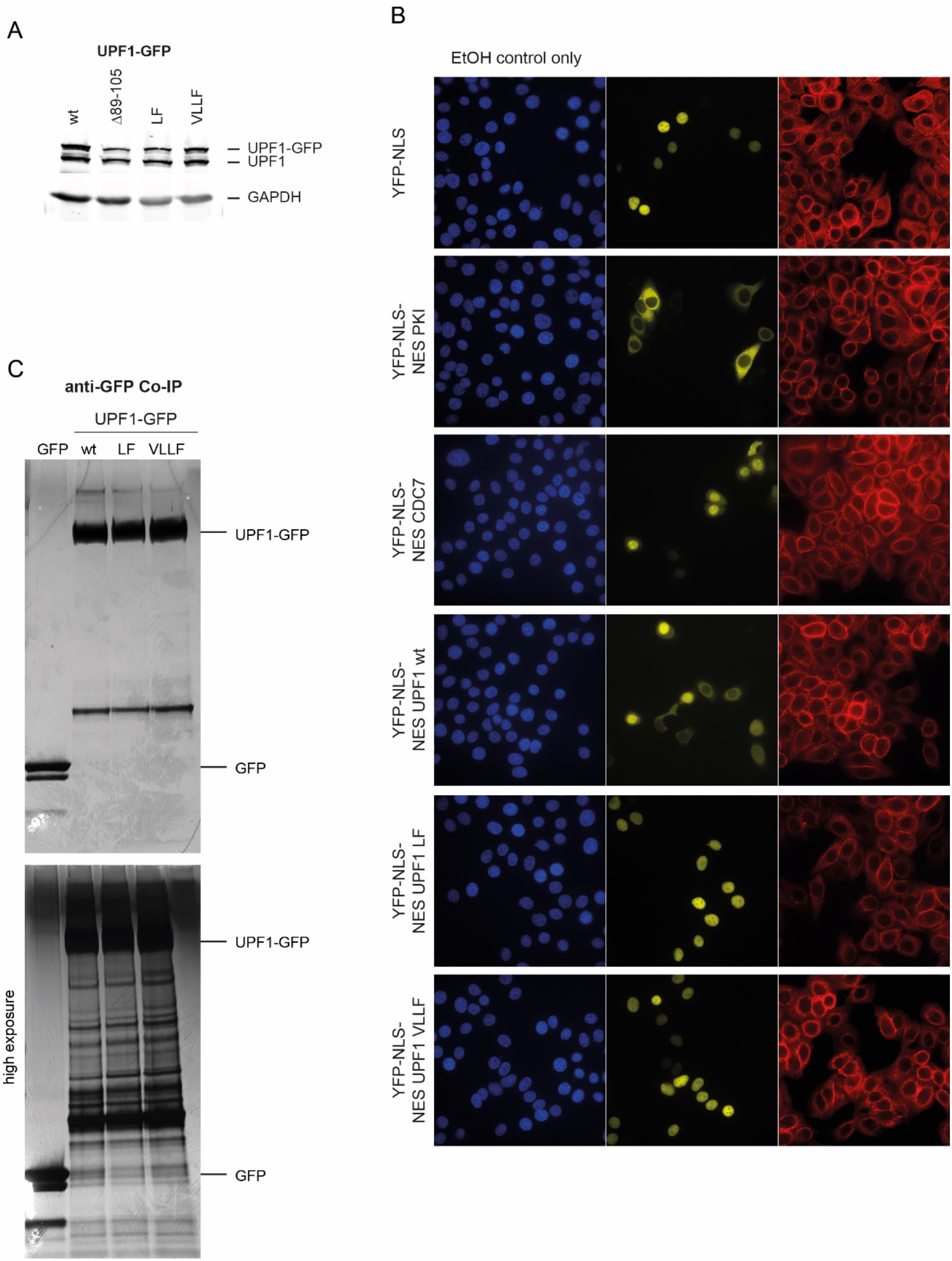
**A**. Immunoblotting to confirm the expression of UPF1-GFP variants used in Fig. 4 (Western blot as in Fig. S1). **B**. Pictures of confocal fluorescence microscope of the blue (DAPI), yellow (YFP-NLS), and red (Phalloidin) channels. The six different constructs are explained in the Results section and in the legend of Fig. 4B. Only the control condition (EtOH control) is shown. **C**. Co-immunoprecipitation experiments from Fig. 5A was analyzed by SDS-PAGE and silver staining.

**Figure S3.**
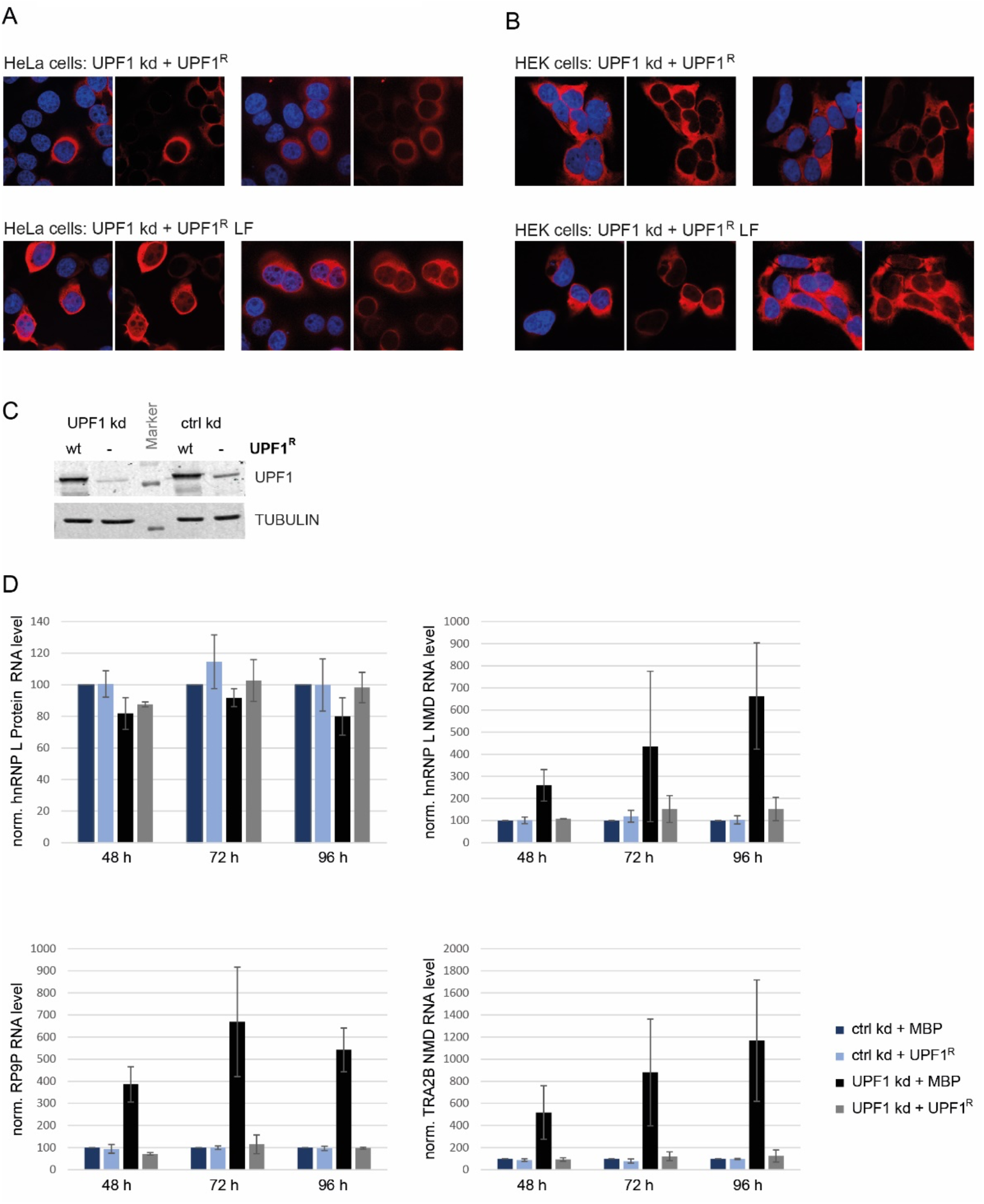
**A**. Immunofluorescence experiments of HeLa cells transiently expressing UPF1 wt or LF mutant in UPF1 kd as in Fig. 5B to investigate the localization of Flag-tagged UPF1 (in red) by confocal microscopy. DAPI was used to stain the nucleus. **B**. Immunofluorescence experiments of HEK cells induced for 96 hours with Dox to express siRNAs against UPF1 (UPF1 kd) or against luciferase (ctrl kd) and RNAi resistant UPF1 wt or LF mutant was analyzed as in A. **C**. UPF1 kd efficacy was assessed by Western blotting of HEK cell lysates 96 hours after addition of Dox as in Fig. 5C. Tubulin served as loading control. **D**. Relative levels of hnRNP L Protein, hnRNP L NMD, RP9P, and TRA2B NMD mRNAs, normalized to actin mRNA, were analyzed by RT-qPCR in HEK cells 48, 72 or 96 hours after addition of Dox. The different HEK cell lines are explained in the Results section and in Fig. 5C. Averages and standard deviations of three independent experiments are shown.

**Table S1.**
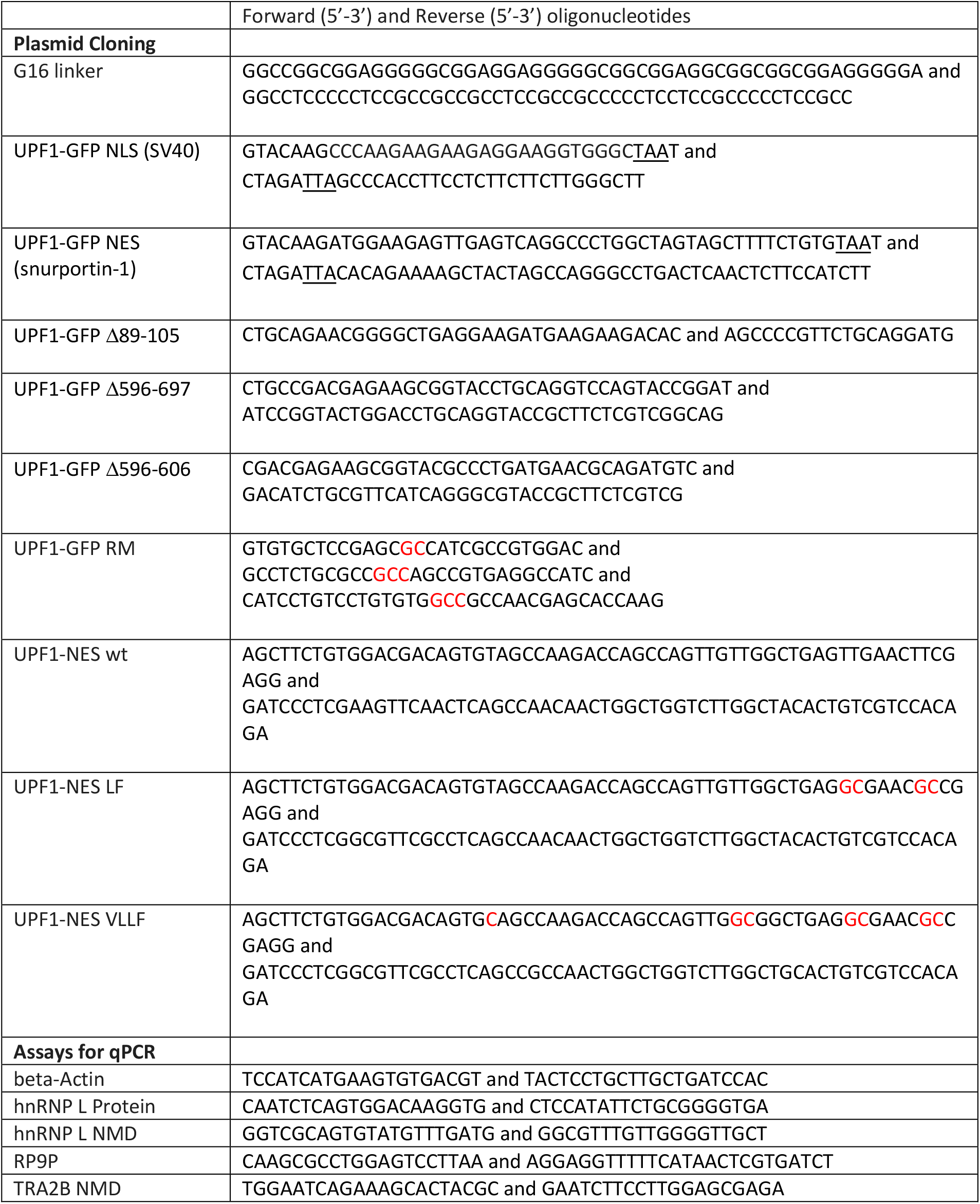
Oligonucleotides (from Microsynth, Switzerland) used in the study. Termination codons are underlined and changes to alanines are marked in red.

## References

Ajamian, L., Abel, K., Rao, S., Vyboh, K., Garcia-de-Gracia, F., Soto-Rifo, R., Kulozik, A.E., Gehring, N.H., and Mouland, A.J. (2015). HIV-1 Recruits UPF1 but Excludes UPF2 to Promote Nucleocytoplasmic Export of the Genomic RNA. Biomolecules 5, 2808–2839. 10.3390/biom5042808.

Azzalin, C.M., and Lingner, J. (2006). The human RNA surveillance factor UPF1 is required for S phase progression and genome stability. Curr Biol 16, 433–439.

Azzalin, C.M., Reichenbach, P., Khoriauli, L., Giulotto, E., and Lingner, J. (2007). Telomeric repeat containing RNA and RNA surveillance factors at mammalian chromosome ends. Science 318, 798–801. 10.1126/science.1147182.

Bhattacharya, A., Czaplinski, K., Trifillis, P., He, F., Jacobson, A., and Peltz, S.W. (2000). Characterization of the biochemical properties of the human Upf1 gene product that is involved in nonsense-mediated mRNA decay. RNA 6, 1226–1235.

Boehm, V., Kueckelmann, S., Gerbracht, J.V., Kallabis, S., Britto-Borges, T., Altmuller, J., Kruger, M., Dieterich, C., and Gehring, N.H. (2021). SMG5-SMG7 authorize nonsense-mediated mRNA decay by enabling SMG6 endonucleolytic activity. Nature communications 12, 3965. 10.1038/s41467-021-24046-3.

Carpenter, A.E., Jones, T.R., Lamprecht, M.R., Clarke, C., Kang, I.H., Friman, O., Guertin, D.A., Chang, J.H., Lindquist, R.A., Moffat, J., et al. (2006). CellProfiler: image analysis software for identifying and quantifying cell phenotypes. Genome Biol 7, R100. 10.1186/gb-2006-7-10-r100.

Chakrabarti, S., Jayachandran, U., Bonneau, F., Fiorini, F., Basquin, C., Domcke, S., Le Hir, H., and Conti, E. (2011). Molecular mechanisms for the RNA-dependent ATPase activity of Upf1 and its regulation by Upf2. Mol Cell 41, 693–703. 10.1016/j.molcel.2011.02.010.

Chamieh, H., Ballut, L., Bonneau, F., and Le Hir, H. (2008). NMD factors UPF2 and UPF3 bridge UPF1 to the exon junction complex and stimulate its RNA helicase activity. Nat Struct Mol Biol 15, 85–93.

Chawla, R., Redon, S., Raftopoulou, C., Wischnewski, H., Gagos, S., and Azzalin, C.M. (2011). Human UPF1 interacts with TPP1 and telomerase and sustains telomere leading-strand replication. EMBO J 30, 4047–4058. 10.1038/emboj.2011.280.

Cheng, Z., Muhlrad, D., Lim, M.K., Parker, R., and Song, H. (2007). Structural and functional insights into the human Upf1 helicase core. EMBO J 26, 253–264.

Cho, H., Park, O.H., Park, J., Ryu, I., Kim, J., Ko, J., and Kim, Y.K. (2015). Glucocorticoid receptor interacts with PNRC2 in a ligand-dependent manner to recruit UPF1 for rapid mRNA degradation. Proc Natl Acad Sci U S A 112, E1540–1549. 10.1073/pnas.1409612112.

Choe, J., Ahn, S.H., and Kim, Y.K. (2014). The mRNP remodeling mediated by UPF1 promotes rapid degradation of replication-dependent histone mRNA. Nucleic Acids Res 42, 9334–9349. 10.1093/nar/gku610.

Contu, L., Balistreri, G., Domanski, M., Uldry, A.C., and Muhlemann, O. (2021). Characterisation of the Semliki Forest Virus-host cell interactome reveals the viral capsid protein as an inhibitor of nonsense-mediated mRNA decay. PLoS Pathog 17, e1009603. 10.1371/journal.ppat.1009603.

Dehghani-Tafti, S., and Sanders, C.M. (2017). DNA substrate recognition and processing by the full-length human UPF1 helicase. Nucleic Acids Res 45, 7354–7366. 10.1093/nar/gkx478.

Durand, S., Franks, T.M., and Lykke-Andersen, J. (2016). Hyperphosphorylation amplifies UPF1 activity to resolve stalls in nonsense-mediated mRNA decay. Nature communications 7, 12434. 10.1038/ncomms12434.

Eberle, A.B., Lykke-Andersen, S., Muhlemann, O., and Jensen, T.H. (2009). SMG6 promotes endonucleolytic cleavage of nonsense mRNA in human cells. Nat Struct Mol Biol 16, 49–55. 10.1038/nsmb.1530.

Elbarbary, R.A., Miyoshi, K., Hedaya, O., Myers, J.R., and Maquat, L.E. (2017). UPF1 helicase promotes TSN-mediated miRNA decay. Genes Dev 31, 1483–1493. 10.1101/gad.303537.117.

Fiorini, F., Bagchi, D., Le Hir, H., and Croquette, V. (2015). Human Upf1 is a highly processive RNA helicase and translocase with RNP remodelling activities. Nature communications 6, 7581. 10.1038/ncomms8581.

Fiorini, F., Boudvillain, M., and Le Hir, H. (2013). Tight intramolecular regulation of the human Upf1 helicase by its N-and C-terminal domains. Nucleic Acids Res 41, 2404–2415. 10.1093/nar/gks1320.

Franks, T.M., Singh, G., and Lykke-Andersen, J. (2010). Upf1 ATPase-dependent mRNP disassembly is required for completion of nonsense-mediated mRNA decay. Cell 143, 938–950. 10.1016/j.cell.2010.11.043.

Gowravaram, M., Bonneau, F., Kanaan, J., Maciej, V.D., Fiorini, F., Raj, S., Croquette, V., Le Hir, H., and Chakrabarti, S. (2018). A conserved structural element in the RNA helicase UPF1 regulates its catalytic activity in an isoform-specific manner. Nucleic Acids Res 46, 2648–2659. 10.1093/nar/gky040.

Hein, M.Y., Hubner, N.C., Poser, I., Cox, J., Nagaraj, N., Toyoda, Y., Gak, I.A., Weisswange, I., Mansfeld, J., Buchholz, F., et al. (2015). A human interactome in three quantitative dimensions organized by stoichiometries and abundances. Cell 163, 712–723. 10.1016/j.cell.2015.09.053.

Hong, D., Park, T., and Jeong, S. (2019). Nuclear UPF1 Is Associated with Chromatin for Transcription-Coupled RNA Surveillance. Mol Cells. 10.14348/molcells.2019.0116.

Huntzinger, E., Kashima, I., Fauser, M., Sauliere, J., and Izaurralde, E. (2008). SMG6 is the catalytic endonuclease that cleaves mRNAs containing nonsense codons in metazoan. RNA 14, 2609–2617. 10.1261/rna.1386208.

Hurt, J.A., Robertson, A.D., and Burge, C.B. (2013). Global analyses of UPF1 binding and function reveal expanded scope of nonsense-mediated mRNA decay. Genome Res 23, 1636–1650. 10.1101/gr.157354.113.

Jankowsky, E. (2011). RNA helicases at work: binding and rearranging. Trends Biochem Sci 36, 19–29. 10.1016/j.tibs.2010.07.008.

Karousis, E.D., Gypas, F., Zavolan, M., and Mühlemann, O. (2021). Nanopore sequencing reveals endogenous NMD-targeted isoforms in human cells. Genome Biology in press. BioRxiv preprint: https://doi.org/10.1101/2021.04.30.442116.

Karousis, E.D., and Mühlemann, O. (2019). Nonsense-Mediated mRNA Decay Begins Where Translation Ends. Cold Spring Harb Perspect Biol 11. 10.1101/cshperspect.a032862.

Kaygun, H., and Marzluff, W.F. (2005). Regulated degradation of replication-dependent histone mRNAs requires both ATR and Upf1. Nat Struct Mol Biol 12, 794–800. 10.1038/nsmb972.

Kazgan, N., Williams, T., Forsberg, L.J., and Brenman, J.E. (2010). Identification of a nuclear export signal in the catalytic subunit of AMP-activated protein kinase. Mol Biol Cell 21, 3433–3442. 10.1091/mbc.E10-04-0347.

Kim, Y.K., Furic, L., Desgroseillers, L., and Maquat, L.E. (2005). Mammalian Staufen1 recruits Upf1 to specific mRNA 3’UTRs so as to elicit mRNA decay. Cell 120, 195–208.

Kim, Y.K., and Maquat, L.E. (2019). UPFront and center in RNA decay: UPF1 in nonsense-mediated mRNA decay and beyond. RNA. 10.1261/rna.070136.118.

Kosugi, S., Hasebe, M., Matsumura, N., Takashima, H., Miyamoto-Sato, E., Tomita, M., and Yanagawa, H. (2009a). Six classes of nuclear localization signals specific to different binding grooves of importin alpha. J Biol Chem 284, 478–485. 10.1074/jbc.M807017200.

Kosugi, S., Hasebe, M., Tomita, M., and Yanagawa, H. (2008). Nuclear export signal consensus sequences defined using a localization-based yeast selection system. Traffic 9, 2053–2062. 10.1111/j.1600-0854.2008.00825.x.

Kosugi, S., Hasebe, M., Tomita, M., and Yanagawa, H. (2009b). Systematic identification of cell cycle-dependent yeast nucleocytoplasmic shuttling proteins by prediction of composite motifs. Proc Natl Acad Sci U S A 106, 10171–10176. 10.1073/pnas.0900604106.

Kudo, N., Wolff, B., Sekimoto, T., Schreiner, E.P., Yoneda, Y., Yanagida, M., Horinouchi, S., and Yoshida, M. (1998). Leptomycin B inhibition of signal-mediated nuclear export by direct binding to CRM1. Exp Cell Res 242, 540–547. 10.1006/excr.1998.4136.

Kurosaki, T., and Maquat, L.E. (2013). Rules that govern UPF1 binding to mRNA 3’ UTRs. Proc Natl Acad Sci U S A 110, 3357–3362. 10.1073/pnas.1219908110.

Kurosaki, T., Popp, M.W., and Maquat, L.E. (2019). Quality and quantity control of gene expression by nonsense-mediated mRNA decay. Nat Rev Mol Cell Biol 20, 406–420. 10.1038/s41580-019-0126-2.

Lee, B.J., Cansizoglu, A.E., Suel, K.E., Louis, T.H., Zhang, Z., and Chook, Y.M. (2006). Rules for nuclear localization sequence recognition by karyopherin beta 2. Cell 126, 543–558. 10.1016/j.cell.2006.05.049.

Lee, S.R., Pratt, G.A., Martinez, F.J., Yeo, G.W., and Lykke-Andersen, J. (2015). Target Discrimination in Nonsense-Mediated mRNA Decay Requires Upf1 ATPase Activity. Mol Cell 59, 413–425. 10.1016/j.molcel.2015.06.036.

Longman, D., Jackson-Jones, K.A., Maslon, M.M., Murphy, L.C., Young, R.S., Stoddart, J.J., Hug, N., Taylor, M.S., Papadopoulos, D.K., and Caceres, J.F. (2020). Identification of a localized nonsense-mediated decay pathway at the endoplasmic reticulum. Genes Dev 34, 1075–1088. 10.1101/gad.338061.120.

Lott, K., and Cingolani, G. (2011). The importin beta binding domain as a master regulator of nucleocytoplasmic transport. Biochim Biophys Acta 1813, 1578–1592. 10.1016/j.bbamcr.2010.10.012.

Lu, J., Wu, T., Zhang, B., Liu, S., Song, W., Qiao, J., and Ruan, H. (2021). Types of nuclear localization signals and mechanisms of protein import into the nucleus. Cell Commun Signal 19, 60. 10.1186/s12964-021-00741-y.

Lykke-Andersen, J., Shu, M.D., and Steitz, J.A. (2000). Human Upf proteins target an mRNA for nonsense-mediated decay when bound downstream of a termination codon. Cell 103, 1121–1131. 10.1016/s0092-8674(00)00214-2.

Lykke-Andersen, S., Chen, Y., Ardal, B.R., Lilje, B., Waage, J., Sandelin, A., and Jensen, T.H. (2014). Human nonsense-mediated RNA decay initiates widely by endonucleolysis and targets snoRNA host genes. Genes Dev 28, 2498–2517. 10.1101/gad.246538.114.

McNally, L.M., Yee, L., and McNally, M.T. (2006). Heterogeneous nuclear ribonucleoprotein H is required for optimal U11 small nuclear ribonucleoprotein binding to a retroviral RNA-processing control element: implications for U12-dependent RNA splicing. J Biol Chem 281, 2478–2488. 10.1074/jbc.M511215200.

Medghalchi, S.M., Frischmeyer, P.A., Mendell, J.T., Kelly, A.G., Lawler, A.M., and Dietz, H.C. (2001). Rent1, a trans-effector of nonsense-mediated mRNA decay, is essential for mammalian embryonic viability. Hum Mol Genet 10, 99–105.

Mendell, J.T., ap Rhys, C.M., and Dietz, H.C. (2002). Separable roles for rent1/hUpf1 in altered splicing and decay of nonsense transcripts. Science 298, 419–422. 10.1126/science.1074428.

Mino, T., Murakawa, Y., Fukao, A., Vandenbon, A., Wessels, H.H., Ori, D., Uehata, T., Tartey, S., Akira, S., Suzuki, Y., et al. (2015). Regnase-1 and Roquin Regulate a Common Element in Inflammatory mRNAs by Spatiotemporally Distinct Mechanisms. Cell 161, 1058–1073. 10.1016/j.cell.2015.04.029.

Ngo, G.H.P., Grimstead, J.W., and Baird, D.M. (2021). UPF1 promotes the formation of R loops to stimulate DNA double-strand break repair. Nature communications 12, 3849. 10.1038/s41467-021-24201-w.

Nicholson, P., Joncourt, R., and Muhlemann, O. (2012). Analysis of nonsense-mediated mRNA decay in mammalian cells. Curr Protoc Cell Biol Chapter 27, Unit27 24. 10.1002/0471143030.cb2704s55.

Nicholson, P., Josi, C., Kurosawa, H., Yamashita, A., and Muhlemann, O. (2014). A novel phosphorylation-independent interaction between SMG6 and UPF1 is essential for human NMD. Nucleic Acids Res 42, 9217–9235. 10.1093/nar/gku645.

Okamura, M., Inose, H., and Masuda, S. (2015). RNA Export through the NPC in Eukaryotes. Genes (Basel) 6, 124–149. 10.3390/genes6010124.

Paillusson, A., Hirschi, N., Vallan, C., Azzalin, C.M., and Muhlemann, O. (2005). A GFP-based reporter system to monitor nonsense-mediated mRNA decay. Nucleic Acids Res 33, e54.

Park, E., and Maquat, L.E. (2013). Staufen-mediated mRNA decay. Wiley Interdiscip Rev RNA 4, 423–435. 10.1002/wrna.1168.

Park, O.H., Park, J., Yu, M., An, H.T., Ko, J., and Kim, Y.K. (2016). Identification and molecular characterization of cellular factors required for glucocorticoid receptor-mediated mRNA decay. Genes Dev 30, 2093–2105. 10.1101/gad.286484.116.

Rahmani, K., and Dean, D.A. (2017). Leptomycin B alters the subcellular distribution of CRM1 (Exportin 1). Biochem Biophys Res Commun 488, 253–258. 10.1016/j.bbrc.2017.04.042.

Rufener, S.C., and Muhlemann, O. (2013). eIF4E-bound mRNPs are substrates for nonsense-mediated mRNA decay in mammalian cells. Nature structural & molecular biology 20, 710–717. 10.1038/nsmb.2576.

Singh, A.K., Choudhury, S.R., De, S., Zhang, J., Kissane, S., Dwivedi, V., Ramanathan, P., Petric, M., Orsini, L., Hebenstreit, D., and Brogna, S. (2019). The RNA helicase UPF1 associates with mRNAs co-transcriptionally and is required for the release of mRNAs from gene loci. eLife 8. 10.7554/eLife.41444.

Singh, G., Pratt, G., Yeo, G.W., and Moore, M.J. (2015). The Clothes Make the mRNA: Past and Present Trends in mRNP Fashion. Annu Rev Biochem. 10.1146/annurev-biochem-080111-092106.

Sun, X., Perlick, H.A., Dietz, H.C., and Maquat, L.E. (1998). A mutated human homologue to yeast Upf1 protein has a dominant-negative effect on the decay of nonsense-containing mRNAs in mammalian cells. Proc Natl Acad Sci U S A 95, 10009–10014.

Szczesny, R.J., Kowalska, K., Klosowska-Kosicka, K., Chlebowski, A., Owczarek, E.P., Warkocki, Z., Kulinski, T.M., Adamska, D., Affek, K., Jedroszkowiak, A., et al. (2018). Versatile approach for functional analysis of human proteins and efficient stable cell line generation using FLP-mediated recombination system. PLoS One 13, e0194887. 10.1371/journal.pone.0194887.

Xu, D., Marquis, K., Pei, J., Fu, S.C., Cagatay, T., Grishin, N.V., and Chook, Y.M. (2015). LocNES: a computational tool for locating classical NESs in CRM1 cargo proteins. Bioinformatics 31, 1357–1365. 10.1093/bioinformatics/btu826.

Yamashita, A., Ohnishi, T., Kashima, I., Taya, Y., and Ohno, S. (2001). Human SMG-1, a novel phosphatidylinositol 3-kinase-related protein kinase, associates with components of the mRNA surveillance complex and is involved in the regulation of nonsense-mediated mRNA decay. Genes Dev 15, 2215–2228.

Zund, D., Gruber, A.R., Zavolan, M., and Muhlemann, O. (2013). Translation-dependent displacement of UPF1 from coding sequences causes its enrichment in 3’ UTRs. Nature structural & molecular biology 20, 936–943. 10.1038/nsmb.2635.

